# Understanding Activity-Stability Tradeoffs in Biocatalysts by Enzyme Proximity Sequencing

**DOI:** 10.1101/2023.02.24.529916

**Authors:** Rosario Vanella, Christoph Küng, Alexandre A. Schoepfer, Vanni Doffini, Jin Ren, Michael A. Nash

## Abstract

Understanding the complex relationships between enzyme sequence, folding stability and catalytic activity is crucial for applications in industry and biomedicine. However, current enzyme assay technologies are limited by an inability to simultaneously resolve both stability and activity phenotypes and to couple these to gene sequences at large scale. Here we developed Enzyme Proximity Sequencing (EP-Seq), a deep mutational scanning method that leverages peroxidase-mediated radical labeling with single cell fidelity to dissect the effects of thousands of mutations on stability and catalytic activity of oxidoreductase enzymes in a single experiment. We used EP-Seq to analyze how 6,399 missense mutations influence folding stability and catalytic activity in a D-amino acid oxidase (DAOx) from *R*.*gracilis*. The resulting datasets demonstrate activity-based constraints that limit folding stability during natural evolution, and identify hotspots distant from the active site as candidates for mutations that improve catalytic activity without sacrificing stability. EP-Seq can be extended to other enzyme classes and provides valuable insights into biophysical principles governing enzyme structure and function.

## Introduction

Soluble proteins produced through natural selection are typically only marginally stable. For enzymes, local flexibility is required at the active site to achieve catalytic activity, however excessive mobility renders them susceptible to denaturation. This tradeoff between activity and stability is still not well understood in protein science^1–4^. Observations on small numbers of homologous sequences have shown how cold-adapted enzymes are typically more active than thermophilic homologs^5–7^, however, competing studies relying on database meta-analysis^8^, experimental data and comparative phylogenetics^9^ have challenged this thermal rate compensation model. The question of how enzymes encode activity and stability, and the interplay between the two over the course of natural evolution or during experimental laboratory directed evolution^10,11^ therefore remains open. For enzyme engineering, these tradeoffs mean that both stability and activity cannot be optimized simultaneously or with equal success. If we could experimentally determine sequence-function relationships describing folding stability and catalytic activity of enzyme variants at large scale, it would enable an understanding of the activity-stability tradeoff, and potentially unlock enhanced enzymes for industrial and biomedical applications.

With the advent of deep mutational scanning (DMS), the effects of large numbers of genetic mutations on protein phenotype can be analyzed using massively parallel methods^12–14^. However, catalytic enzymes are challenging for DMS studies because there exist very few massively scalable screening methods which successfully link genotype and phenotype. Typically, enzyme fitness is coupled to host cell survival using growth-based selection. Alternatively, microdroplet methods allow single clones to be analyzed using colorimetric assays followed by droplet sorting and sequencing. Enrichment of variants from pre-to post-selected pools allows tabulation of phenotypic scores and provides insights into mutational fitness landscapes^15–21^. In nearly all prior implementations of DMS on enzymes, however, fitness scores comprise a conflation of expression level (i.e. enzyme abundance) and catalytic activity. Klesmith and colleagues showed how enzyme fitness scores determined through growth-based selection could be combined with solubility scores from independent assays to reveal evolutionary origins of stability activity trade-offs^22^. Markin *et al*. further presented a microfluidic enzyme expression platform which decoupled the catalytic properties of each variant from their expression levels^23^. While very powerful, each platform has associated limitations in terms of throughput, ease-of-use, compatible chemistries, and robustness.

Here we report EP-Seq, a novel DMS-based method that combines enzyme proximity labeling with next-generation DNA sequencing (NGS) technology to assay both expression level and catalytic activity phenotypes of thousands of variant enzymes from a cellular pool in a single experiment. EP-Seq leverages features of yeast surface display^24,25^ to measure the expression levels of each enzyme variant *via* a pooled cell sorting-sequencing experiment. In parallel, phenoxyl radical-based^26–28^ cell surface proximity labeling links enzyme activity to a fluorescent signal, which is then quantified by sorting and sequencing. We used EP-Seq to study a nearly comprehensive site saturation mutational library of D-amino acid oxidase from the yeast *Rhodotorula gracilis*, a model flavoprotein with industrial and therapeutic applications^29–34^. Downstream computational analysis of EP-Seq data reveals rich biophysical insights into the enzyme by quantifying fitness propensities of substituted residues, identifying protein regions where catalytic activity served as an evolutionary constraint on folding stability, and making predictions of catalysis-enhancing mutations that maintain folding stability.

## Results and discussion

### Workflow overview

An overview of the EP-Seq workflow is shown in Figure 1. In one branch of the experiment (Fig. 1, left), we analyze the expression levels of thousands of variants in parallel by displaying a variant library on yeast, staining the expressed proteins with fluorescent antibodies, and sorting the cells into 4 bins using fluorescence activated cell sorting (FACS). We then use NGS to sequence the variants in each bin, and convert the NGS reads into expression fitness scores using a custom computational pipeline. There is strong evidence that the level at which yeasts secrete and display a given protein sequence is correlated with its folding stability^22,25^. Destabilizing mutations can activate the yeast quality control system, exposing variants to proteases in the secretory pathway which degrade unstable sequences and lower expression levels^35^. The expression level of a variant enzyme analyzed through FACS and DMS can therefore serve as a proxy for folding stability. In this study, we use the term “folding stability” to describe the impact of mutations on the overall cellular stability of the target protein. This primarily relates to structural and thermodynamic stability, but can also include other factors like mRNA stability, efficiency of translation and secretion, and susceptibility to protease degradation, all of which contribute to changes in the protein’s expression level.

**Figure 1.**
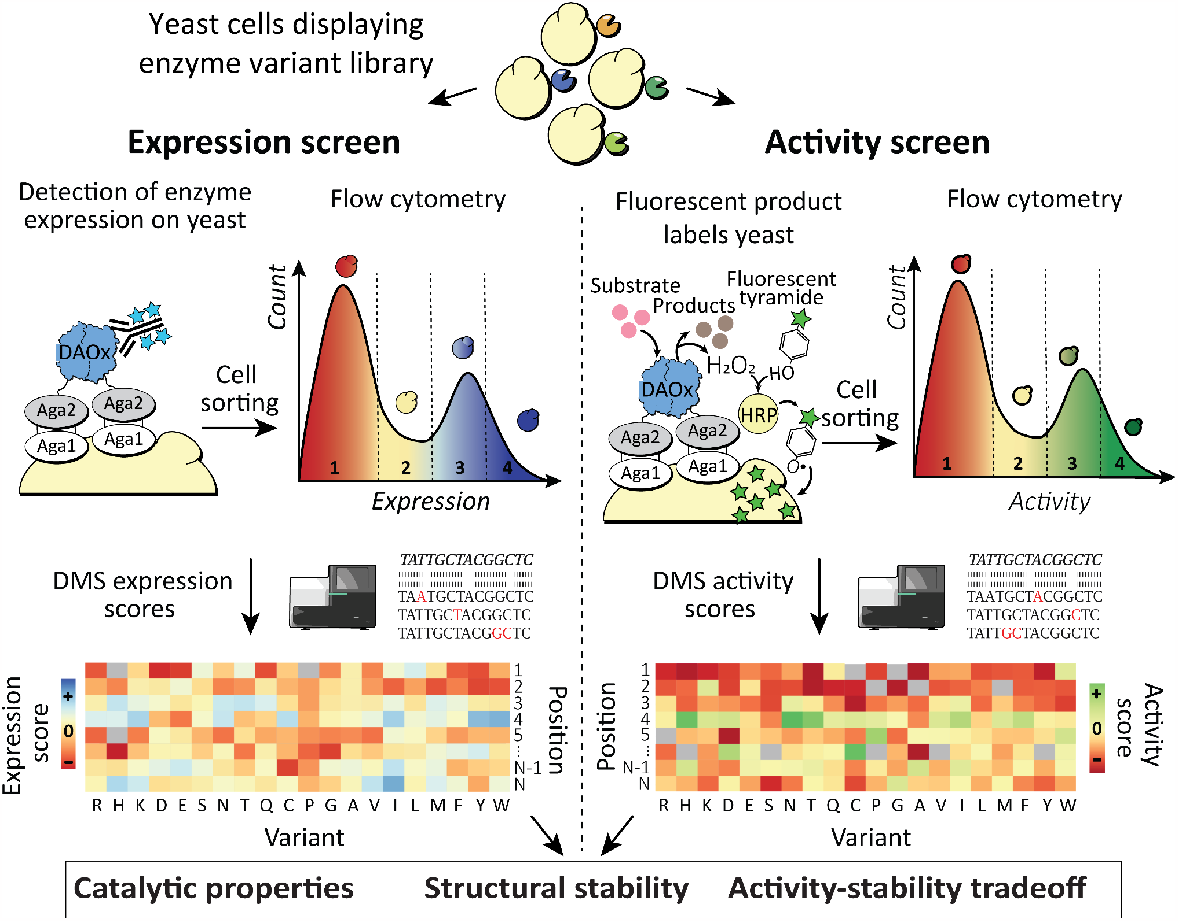
Schematic depicting the EP-Seq workflow. (Top) A pooled library of enzyme variants is displayed on yeast. (Left) The cell population is sorted into bins based on the expression level of the displayed enzyme. (Right) The pooled variant library is assayed for DAOx activity using a cascade peroxidase-mediated proximity labeling reaction with single cell fidelity and sorted into bins. The genetic composition of cells in the sorted bins is quantified via high-throughput sequencing and the distribution of each variant along the expression and activity axes is converted into a fitness score. Joint analysis of the two independent datasets provides insights into the effects of mutations on structural stability and activity of the enzyme.

In a parallel branch of the experiment (Fig. 1, right), the oxidase activity of enzyme variants is assayed at large scale in a pooled format using a horseradish peroxidase (HRP)-mediated phenoxyl radical coupling reaction at the yeast surface. Similar proximity labeling reaction schemes relying on HRP or ascorbate peroxidase-2 (APEX2) have been used in quantitative proteomics^36^, proximity labeling in live cells^37–40^, electron microscopy labelling^41,42^ and signal amplification in biosensors^43,44^. In EP-Seq, the short half-life of phenoxyl radicals limits the labeling reaction to the surface of the cell that generates H_2_O_2_, affording an artificial quasi single-cell reaction compartment by means of a reaction-diffusion limitation. The pooled cells displaying the variant library stained in this fashion are sorted into bins based on fluorescent intensity, and sequenced by NGS. Finally, the large datasets obtained in the two parallel screening protocols are combined, cross-referenced and studied to reveal biophysical insights into protein sequence, stability, and function.

### Quantifying DAOx stability and catalytic activity fitness with EP-Seq scores

We applied the workflow described above to study the 80 kDa homodimeric flavin adenine dinucleotide (FAD) dependent DAOx from *Rhodotorula gracilis* (*Rg*), which promiscuously catalyzes deamination of D-amino acids to imino acids, generating H_2_O_2_ as a byproduct^34^. DAOx has attracted attention for applications in both industry and biomedicine, for example, in resolving racemic amino acid mixtures, producing antibiotics^31^ or as a proposed cancer therapy via reactive oxygen species^29,30,45^. We first optimized the functional expression and display of the wild type DAOx (WT DAOx) fused with the Aga2 anchor protein and established the tyramide-based proximity labeling assay to detect the activity of each displayed enzyme with single-cell fidelity (Supplementary Note 1). Next, we constructed a library for DMS analysis through site saturation mutagenesis over the entire coding region of DAOx and assigned 15 nucleotides unique molecular identifier (UMIs) to each DAOx variant of the library (Supplementary Note 2).

We investigated the effects of single amino acid substitutions on DAOx expression and display at the yeast surface (Fig. 2, left). Following induction (48 h, 20°C, pH 7), we stained the C-terminal His-tag of the DAOx variant library with primary and fluorescent secondary antibodies. We sorted the library into 4 bins based on expression level, where the non-expressing bin was set using a negative control cell population incubated with only the secondary antibody. The remaining yeasts were sorted into three sub-populations containing equal percentages of expressing cells (Fig. 2*A*). After sorting, we extracted plasmid DNA from each sorted cell population, PCR amplified the regions corresponding to the UMIs, and sent the amplicons for single end (SE100) Illumina sequencing on a NovaSeq 6000. The number of reads per sample was on average 25-fold higher than the number of cells sorted into the corresponding FACS bin (Table S2). We filtered the UMI sequences by read quality (Phred score >= Q20) and expected size (15 nucleotides) before assigning them to the corresponding DAOx variants using the look-up table. We converted the number of reads per variant into number of cells (Methods, Equation 1) and calculated a final expression score (Exp) for each variant (Methods, Equation 2). The fitness score per variant was then calculated as log_2_(β_v_/β_*wt*_) where β_v_ was the weighted mean expression score of the variant enzyme and β_*wt*_ was the score of WT DAOx (Methods, Equation 3).

**Figure 2.**
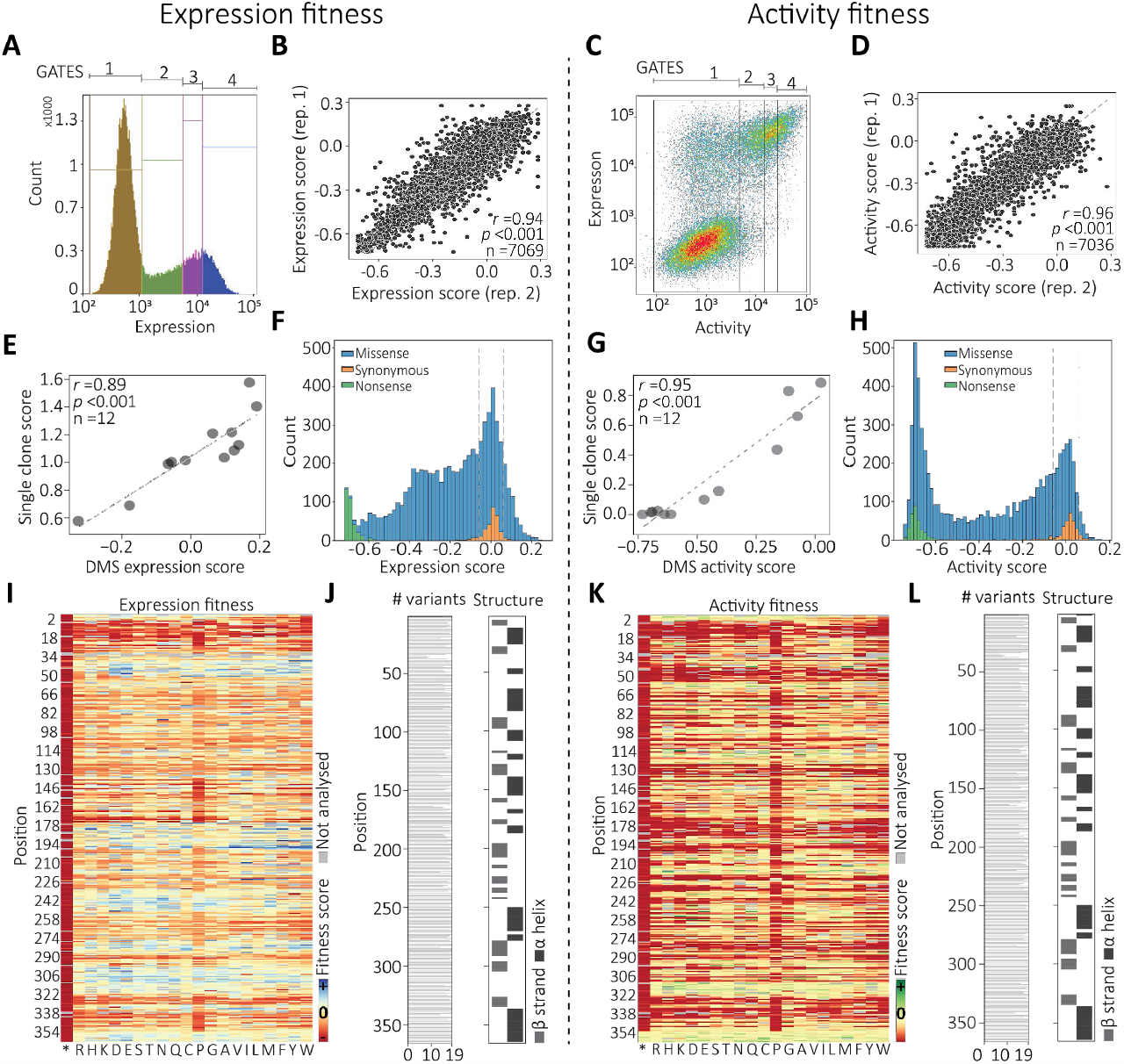
Deep mutational scanning of DAOx expression and catalytic activity by EP-Seq. A) Sorting gates for analysis of display levels. B) Linear correlation between expression scores calculated from two biological replicates. C) Sorting gates for catalytic activity screening. D) Linear correlation between activity scores calculated from two biological replicates. E) Correlation between variant surface display level measured in monogenic yeast culture vs. DMS expression fitness analyzed by EP-Seq for 12 DAOx single mutant variants. F) Distribution of expression fitness effects measured by EP-Seq. Dashed lines represent the range of fitness score for synonymous variants. G) Correlation between variant activity level measured in monogenic yeast culture via peroxidase assay (Amplex Red) vs. DMS activity fitness analyzed by EP-Seq for 12 DAOx single mutant variants. H) Distribution of activity fitness effects measured by EP-Seq. Dashed lines represent the range of fitness score for synonymous variants. I) Expression fitness scores for each variant represented as a heatmap. J) Number of variants analyzed per position in the expression dataset, and secondary structure classification per position (PDB 1C0P). K) Activity fitness scores for each variant obtained by EP-Seq represented as a heatmap. L) Number of variants analyzed per position in the activity dataset and secondary structure classification per position (PDB 1C0P). Links to interactive and color blind accessible heatmaps can be found in the “Data availability statement” section of the manuscript.

To analyze DAOx catalytic activity in a massively parallel fashion, we used a reaction cascade that converts DAOx enzymatic activity into a fluorescent label on the cell wall. This approach is related to prior work from our lab and others on enzyme-mediated polymerization and peroxidase-based proximity labeling^14,27,28,46,47^. We set the low fluorescence gate to include entirely the population of not displaying cells or displaying inactive variants (Fig. 2*C*, Gate 1). The remaining cells were equally divided into three populations corresponding to increasing levels of tyramide-488 signal (Fig. 2*C*). We determined the genetic sequences and their relative abundance in each bin through Illumina sequencing as described above, and identified the corresponding mutant enzyme sequences using the look-up table. The activity score (Act) per variant was calculated (Methods, Equation 2) and the activity fitness score was computed by using as reference the score of the wild type DAOx (Methods, Equation 3). In both the expression and activity screens, we determined a consensus score for each mutant (Methods, Equation 4) and we assessed the reproducibility of the DMS workflow by calculating Pearson’s *r* value for a linear regression of the fitness scores measured in two biological replicates and represented by at least 10 total cells. For the expression assay replicates, the *r* value was 0.94 (n=7,069; *p*<0.001), and for the activity assay replicates, it was 0.96 (n=7,036; *p*<0.001)(Fig. 2*B, D*).

### Validation of EP-Seq scores

To validate the DMS fitness scores of single clones, we randomly selected 12 variants and tested them individually using bulk expression and activity assays. The expression levels of the variants measured individually using yeast display and flow cytometry strongly correlated with expression fitness scores obtained from DMS (*r*=0.89, *p*<0.001) (Fig. 2*E*). The activity levels of the same 12 single mutant variants were next measured using an Amplex Red assay, and the initial reaction rate was measured for the mutants and the wild type enzymes. The single clone score for each mutant was obtained by dividing initial reaction rate of the mutant by that of wild type. All mutants tested showed activity levels consistent with those obtained from EP-Seq (*r*=0. 96, *p*<0.001) (Fig. 2*G*).

To further validate the observed scores, we plotted a histogram to visualize the expression (Fig. 2*F*) and activity (Fig. 2*H*) fitness scores of all single nucleotide variants in the DAOx gene and color coded them to distinguish between missense, nonsense (i.e. stop codon), and synonymous mutations. We found that the distribution of missense variants scores for both expression (n=6,434 variants) and activity (n=6, 404 variants) fitness were broadly distributed (Exp min=-0.72, Exp max=0.25, Act min=-0.75, Act max= 0.16). Fitness score distributions were on average offset towards negative fitness values, indicating a generally deleterious effect of mutation on both expression and activity (median Exp = -0. 15, median Act = -0. 29, Fig. 2*F*,*H*). The distribution of fitness scores containing synonymous mutations in the expression screen was centered at 0.00 ± 0.057 (n=301 variants), while in the activity screen it was centered at 0.00 ± 0. 056 (n=300 variants). This range of fitness values defined a neutral fitness range of the assay. Variants whose scores fell in this range were expected to have fitness comparable to that of WT. Additionally, mutants with nonsense mutations (i.e. stop codons) had strong negative fitness on both expression and activity, with average expression score of -0.69 ± 0.04 (n=334) (Fig. 2*F*) and average activity score of -0.67±0.12 (n=332). The higher activity scores and standard deviation of nonsense variants compared to expression scores of the same variants are due to the effect of stop codon at C-terminal positions of the enzyme (position>350) preserving the formation of a fully folded and functional DAOx enzyme (Fig. *2K*).

As further validation, we calculated the average expression and activity scores per position (n=364) and mapped the values onto the DAOx structure (PDB 1C0P) (Fig. S4, *A,B*). We tested for EP-Seq score correlation with a protein stability prediction algorithm (FOLDX)^48^ (Methods, Equation 5). We used FOLDX to calculate the mean ΔΔG score per position of the protein (n=354) and compared them to expression fitness scores obtained from the left branch of EP-Seq. We found strong correlations between EP-Seq expression scores and predicted ΔΔG values (Exp rho=-0.51 *p*<0.001, Fig. S4*C*), supporting the use of yeast surface display for evaluating the effects of mutations on thermodynamic folding stability. We further compared the predicted ΔΔG values to EP-Seq catalytic activity scores per position of DAOx and again found significant correlation (Act rho=-0. 59 *p*<0.001, Fig. S4*D*). This result supports the idea that a baseline level of structural stability is required for an enzyme to be catalytically active, and that part of the EP-Seq activity score is attributable to effects of mutations on the structural stability (and therefore expression) of DAOx.

### Mutability landscapes of DAOx stability and catalytic activity

We visualized the mutational effects of the 6, 768 (6,434 missense, 334 nonsense) and 6, 736 (6,404 missense, 332 nonsense) mutations in the expression and activity screens (respectively) as fitness heatmaps (Figures 2 I,K). The numbers of missense variants corresponded to 93% and 92. 5% (respectively) of all possible single amino acid substitutions in DAOx (Fig. 2J,L). The expression heatmap (Fig. 2*I*) reveals patterns of higher and lower tolerance for mutation along the DAOx sequence, discussed in detail below. The N-terminal region (residues 8-32) was found to be highly intolerant to amino acid substitutions. This suggests it plays a role in folding stability and could act as an N-terminal intra-molecular chaperone^49^. The Rossmann fold is highly conserved in this region, making contact with the FAD cofactor, stabilizing structure and maintaining function of the enzyme^34^.

### Biophysical properties shape both expression and activity landscapes

We evaluated the effect of mutant residue identity on both expression and activity of DAOx using data from 6, 399 single missense variants for which both EP-Seq expression and activity scores were available (Fig. 3*A, B*, Table S4, Supplementary Note 3). We observed that the introduction of proline had the most pronounced negative impact on both expression and activity, particularly when located in the α helix and β sheet regions of DAOx (Fig. S5*A,B*, Supplementary Note 4). The introduction of cysteine, neutral polar and non-polar residues (excluding glycine) had a positive impact on the studied properties, while the introduction of charged residues (excluding histidine) resulted in a lower activity score than average. Residues with negative charges, glutamic and aspartic acids, exhibited an overall neutral to positive effect on the expression score compared to the average effect of all substituted residues. Finally, among hydrophobic residues, introduction of bulky tryptophan impaired both expression and activity fitness (Fig. 3*A*, Table S4). As a general trend we also observed that DAOx linker regions had greater tolerance to mutations in both expression and activity screens compared to structured regions of the same enzyme (Fig. S5*C,D*, Supplementary Note 4). We next analyzed expression and activity fitness as a function of the substituted wild type amino acids (Fig. S5*E,F*, Supplementary Note 5) and found the substitution of hydrophobic core aromatic residues together with valine, which is one of the most abundant residues in DAOx (n=27), to have the highest negative impact on the expression and activity of the enzyme. On the contrary, mutation of polar and charged amino acids was associated with on average higher scores than the other classes of residues (Fig. S5*E*,F, Supplementary Note 5).

**Figure 3.**
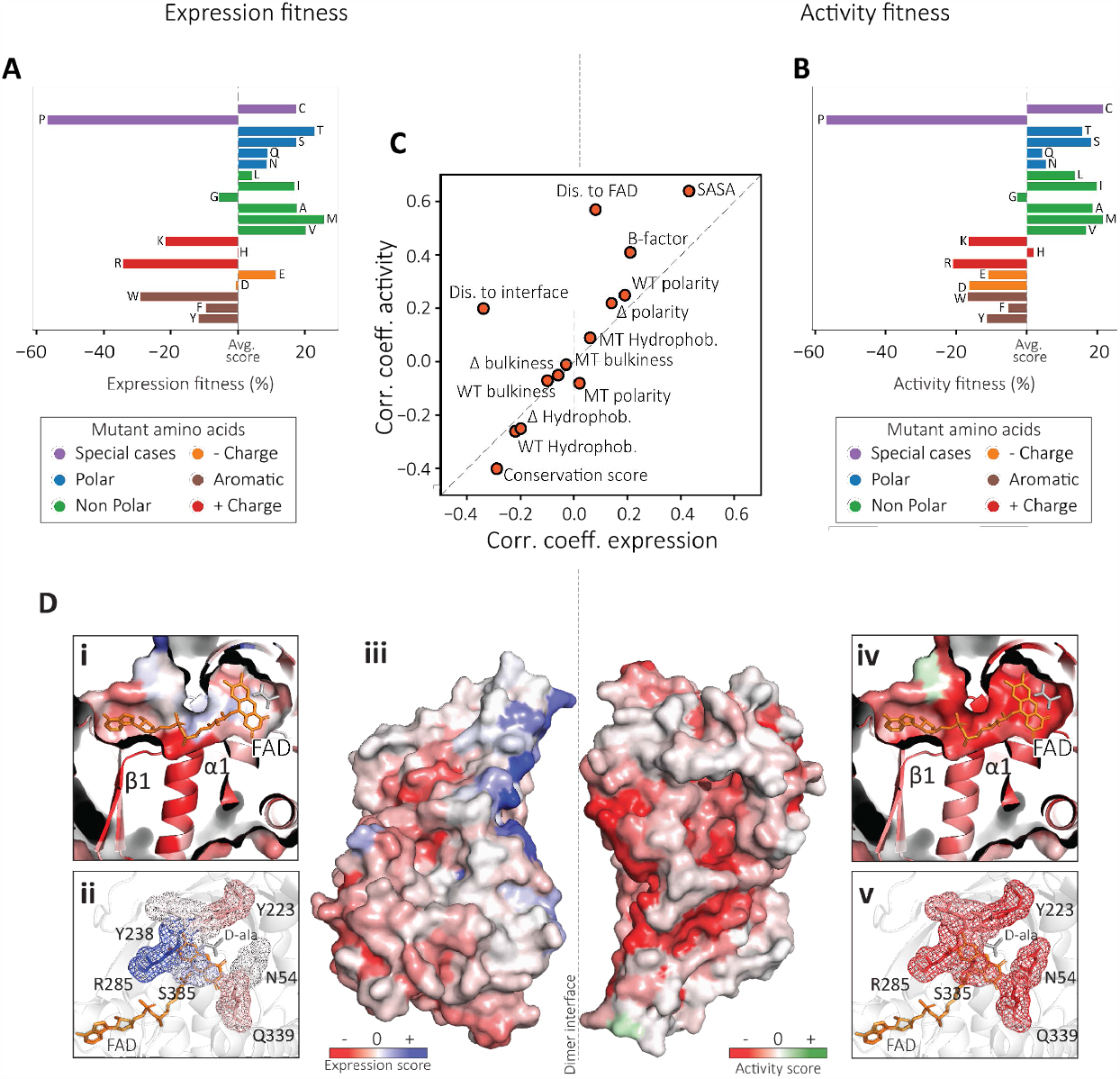
EP-Seq reveals biophysical properties that differentially influence expression and activity. A) Relative impact of mutant residue identity on DAOx expression. B) Relative impact of mutant residue identity on the DAOx enzymatic activity. C) Plot of Pearson correlation coefficients (*r*) calculated for expression (x-axis) or activity (y-axis) fitness scores with respect to several biophysical properties of mutated sites in DAOx. D) DAOx structure colored by expression or fitness scores: (i) Residues surrounding FAD cofactor colored by expression score; (ii) Residues contacting the substrate D-alanine colored by expression score; (iii) Expression (left) or activity (right) scores used to color the surface of DAOx monomers. The monomers are shown separated from each other to improve visibility of the interface; (iv) Residues surrounding FAD cofactor colored by activity score; (v) Residues contacting the substrate D-alanine colored by activity score.

Next, we assigned residue specific scores for hydrophobicity, bulkiness and polarity of wild type and mutant amino acids ^50^ to each of the 6,399 single missense mutations registered by both assays and calculated a Pearson linear correlation coefficient *r* between the values of each feature and the experimental fitness scores (Fig. 3*C*, Fig. S6*A*). We observed no or very low linear correlation between the experimental scores and properties of mutant residues (Mut. hydrophobicity, Mut. polarity, Mut. bulkiness, Fig. S6*A*). Consistent with our previous observations, the hydrophobicity of wild type residues negatively correlated with both expression (*r*=-0.22, *p*<0.001) and activity scores (*r*=-0.26, *p*<0.001), suggesting low tolerance for mutations in the protein core (WT, Δ hydrophobicity, Fig. 3*C*, Fig. S6*A*). We found moderate negative correlation of both expression and activity datasets when considering the size of wild type amino acids or the size difference between wild type and mutant residue side chains (Fig. 3*C*, Fig. S6*A*, WT and Δ bulkiness). Finally, a positive correlation of expression (*r*=0.19, *p*=<0.001) and activity scores (*r*=0.25, *p*=<0.001) with the polarity of wild type amino acids indicated higher tolerance for mutation of polar residues, typically found on the surface of the protein (Fig. 3*C*, Fig. S6*A*, WT and Δ polarity).

### Stability activity trade-offs along natural evolution of DAOx

We next sought to determine whether our expression and activity datasets could be used to identify regions of stability-activity tradeoffs. We analyzed average expression and activity scores of DAOx at each position (n_Epx_=n_Act_=360) in relation to biophysical properties of the mutated sites (Fig. 3*C*) and observed several trends. For example, both expression and activity fitness scores were positively correlated with solvent accessible surface area (SASA) of the mutated site (Expression: *r*=0.43, *p*<0.001; Activity: *r*=0.64, *p*<0.001). The temperature factor (B-factor) was similarly positively correlated with both fitness scores (Expression: *r*=0.21, *p*=<0.001; Activity: *r*=0.41, *p*<0.001) indicating higher mutational tolerance at sites located at the protein surface and at sites with high structural mobility (Fig. 3*C*, Fig. S6*B*,C).

We analyzed fitness scores of mutated sites with respect to their distance from the FAD cofactor (Fig. 3*C*, Dis. to FAD). We found that mutations near FAD often negatively impacted the catalytic activity of the enzyme, as indicated by the positive correlation between the activity scores and the distance of the mutated residue from FAD (*r*=0.57, *p*<0.001, Fig. S6*D* bottom). In contrast, expression fitness scores were not significantly correlated with distance of the mutated residue to FAD (*r*=0. 08, *p*=0.11, Fig. S6*D* top), suggesting that structural stability was insensitive to this parameter. We color-coded the 3D structure of DAOx according to activity and expression scores in the vicinity of the FAD cofactor and in the active site (*Fig 3D i, ii, iv, v*), where we found the largest differences between the activity and expression scores. Mutations in close proximity to FAD (distance<4 Å) greatly impaired the catalytic activity of the enzyme (avg. Act=-0. 557,-66%,n=387; avg. Act all=-0.336, n=6,399; Fig. 3*DIV*) while at the same positions the expression scores showed the opposite trend. This revealed how mutations at the catalytic site tended to harm activity but improve stability (avg. Exp=-0.149, n=387, +17%, avg. Exp all=-0.180,n=6,399; Fig. 3DI), supporting a well-documented phenomenon on the thermodynamic price paid by an enzyme to remain catalytically active under conditions of functional selection ^4^.

A similar behavior was found for residues known to coordinate the substrate D-alanine^34^. While mutating these residues impaired enzyme activity (avg. Act= -0. 457, -36%, n=111; avg. Act all =-0.336, n=6399; Fig. 3*DV*), it tended to improve stability (avg. Exp=-0.06, +60%, n=111; avg. Exp all= -0. 180, n=6399; Fig. 3*DII*). Among the residues interacting with FAD, the loop between strand β1 and helix α1 is highly conserved among Rossmann folds (GSGVIGL, positions: 11-17) and is characterized by negative fitness scores in both datasets (avg. Exp =-0.479, avg. Act=-0.639) (Fig. 3*DI,IV*). This suggests both a functional and structural role of FAD in the overall fitness of the enzyme. These results suggest DAOx folding is facilitated by interactions with FAD, which is also essential for its catalytic activity. The strong binding affinity between DAOx and FAD (K_D_=20 nM) and the low abundance of apo-enzyme further support these observations ^51,52^.

### Dimerization stabilizes marginally stable but functional DAOx monomers

DAOx is active as a homodimer in its native state and dimerization is thought to be required for activity^52,53^. This motivated a detailed analysis of mutations in close proximity to the dimer interface, and whether they could significantly change folding stability and catalytic activity of the enzyme. We calculated the distance between each residue and the closest residue found at the dimer interface, and assigned this distance value to each single missense mutation found in both the expression and activity screens. We then calculated a linear correlation coefficient between the experimental fitness scores and the distance values. We found that the activity fitness data showed a positive correlation with the distance of the mutated site from the dimer interface (*r*=0. 20, *p*<0.001, Fig. 3*C*, Dis. to interface, Fig. S6*E* bottom), while the expression scores correlated negatively with it (*r*=-0.34, *p*=<0.001, Fig. 3*C*, Dis. to interface, Fig. S6*E* top). On average, sites located closer to the dimer interface were more tolerant to mutation in the expression screen, but less so for the activity screen. This reflects a scenario where the enzyme tolerates some amount of instability in order to remain as an active dimer, analogous to the effects at the catalytic site. We visualized these effects on the surface of the 3D structure of the DAOx dimer (Fig. 3D iii). Mutations located near the dimer interface (distance <5 Å) impaired the catalytic activity of the DAOx enzyme as indicated by the dominant red color of the structure (avg. Act=-0. 363, -7%, n=1,298, avg. Act all=-0. 336, n=6,399; Fig. 3D iii, right). The same mutations showed overall positive effects on the expression and stability of the DAOx structure (avg. Exp=-0. 07, +61%, n=1298; avg. Exp all=-0. 180, n=6,399; Fig. 3Diii, left). We attributed this behavior to a strong dependency of DAOx catalytic activity on dimerization^52,53^. Apparently, dimerization along with the conformation of the catalytic site provide functional constraints during evolution of this enzyme, suggesting the evolution of a more stable globular and monomeric form of DAOx might have been impeded during functional selection for catalytic activity.

### EP-Seq expression scores combined with sequence conservation reveal functional sites in DAOx

Sequence conservation scores derived from multisequence alignment reflect constraints imposed on the protein of interest as a result of natural selection. Over evolutionary time, selection tends to maintain both folding stability and catalytic activity^54,55^. We hypothesized that combining conservation scores with our stability and activity datasets could provide insights into functional sites in DAOx. We compared the experimental scores with conservation scores (CONS) obtained by aligning the DAOx sequence with five evolutionarily related D-amino acid oxidase sequences (Fig. S7). Both expression and activity fitness score datasets showed negative correlation with conservation scores (Exp *r*=-29, *p*=2.3e-08; Act *r*=-0.40, *p<0*.*001*, Fig. 3C, Fig. S6*E*), generally indicating higher tolerance for mutations at less conserved protein sites. We identified 41 positions with CONS=100 and analyzed the average expression score per position as a function of the distance of the mutated site from the catalytic center of DAOx. We observed a significant negative correlation of the expression scores with the distance of the mutation from the catalytic site of the enzyme (*r*=-0. 54, *p<0*.*001*, Fig. 4*A*). Among the sites with conservation score 100, we then isolated the positions associated with positive expression score (n=9) and visualized them on the 3D structure of DAOx (Fig. 4*B*). Three of the positions analyzed (A51, G199, R285) found within 5Å of the active site are reported to be directly involved in catalysis through stabilization of the substrate D-alanine (R285) or of the isoalloxazine ring of FAD cofactor (A51, G199)^34^. Conserved residues R198, G199, Q200, G283, and the two prolines in position 196 and 286, are part of two antiparallel strands β8 and β13 suggesting a role of the resulting β sheet in structuring the catalytic pocket and coordinating the side chain of the Arginine 285. Finally, tryptophan W243 located at the surface of monomeric DAOx plays an important role in the dimerization. In agreement with the previous observations, its mutation has a direct impact on the catalytic efficiency of the DAOx^56^ (Fig. 4*B*).

**Figure 4.**
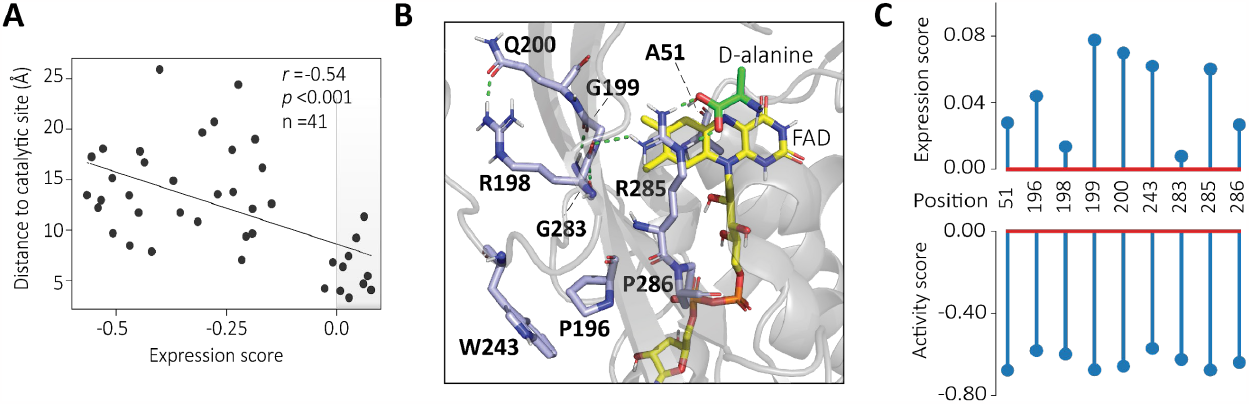
EP-Seq and sequence conservation analysis reveal functional sites of DAOx. A) The distance to the catalytic site for positions that are evolutionarily conserved (conservation score, CONS = 100) were analyzed as a function of EP-Seq expression scores (average per position). The nine highly conserved residues associated with the most positive expression scores (light gray right corner) are visualized in B. B) Structural details of DAOx residues with CONS = 100 and expression score > 0. H-bonds are shown in green. The FAD cofactor and D-alanine carbon backbones are colored in yellow and green, respectively. DAOx residue carbon backbones are colored in light blue with colored elements: Hydrogen (white), Nitrogen (dark blue), Oxygen (Red), Phosphorus (Orange). C) Expression and activity scores for each of the conserved positions linked to positive expression scores.

### Activity enhancing hotspots are globally encoded

We finally used EP-Seq to identify regions of the enzyme where mutations were associated with improved catalytic activity. We generated a two-dimensional scatterplot showing the average activity and expression fitness per position as Y and X coordinates (Fig. 5*A*), and assigned a color at each point based on the distance of the mutated site to the enzyme’s catalytic pocket. The observed trends in this depiction demonstrate how the two properties are related. Variants that are found to be catalytically active must possess at least a minimum level of structural stability in order to be correctly folded, secreted and displayed. This fact is demonstrated by a lack of positions in the upper left of the plot. Only 1 datapoint (∽0.3%) was found with positive activity fitness and negative expression fitness (Fig. 5*A*, top left). Additionally, we found 35 positions (∽10%) exhibiting positive expression fitness with impaired catalytic activity (activity fitness <-0.1) (Fig. 5*A*, lower right). These positions tended on average to be closer to the catalytic site (avg. distance=14.6 Å; avg. distance all=17.4 Å), indicated by the dark blue color in the lower right of the plot. This demonstrates stability-function tradeoffs at play in this enzyme, where more stably expressed variants could be found by mutating crucial catalytic residues at the cost of losing activity. Finally, many mutations which moderately altered the expression of the enzyme resulted in corresponding changes in the catalytic activity (Fig. 5*A*, diagonal). A limited number of positions were tolerant to mutations for both assayed properties (n=23) (Exp>-0.05; Act>-0.05) and showed either simultaneous improvement of both properties or an increase in activity due to higher expression levels on the surface of the yeast cells. These tended to be found more distant from the catalytic site (avg. distance 28.2 Å; avg. distance all=17.4 Å). To further support these observations, we have included a scatter plot to illustrate the relationship between activity and expression fitness for all the single mutant variants included in our work (Fig. S8*A*).

**Figure 5.**
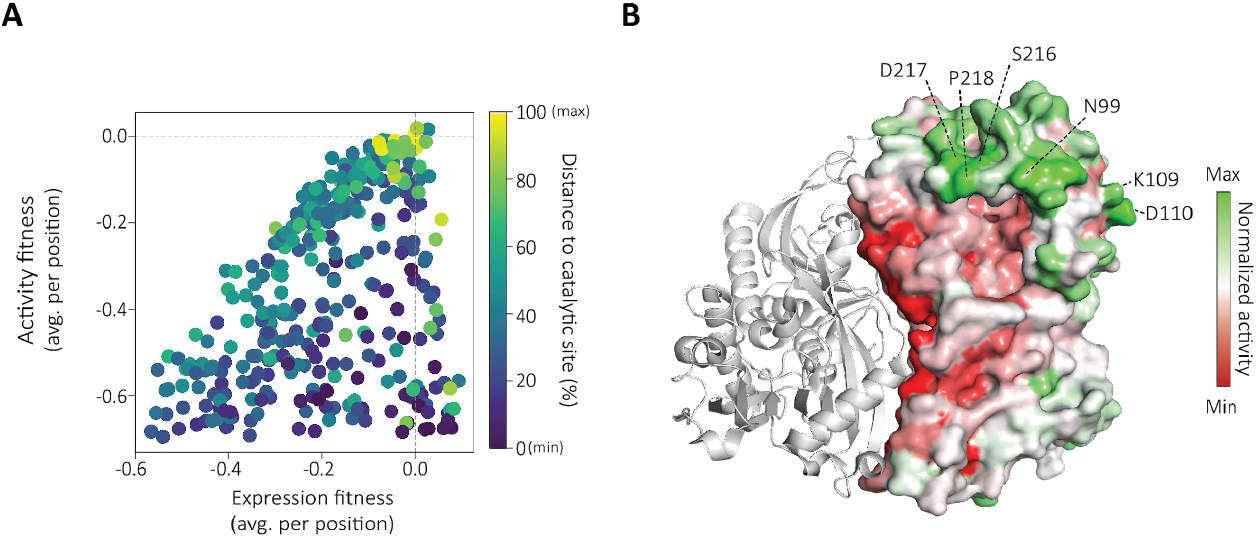
EP-Seq predicts activity enhancing mutations in DAOx. (A) Average expression and activity fitness scores per position in the DAOx enzyme. The coloring indicates distance to the active site. (B) Normalized activity values are depicted on the 3D structure of a DAOx monomer (right). Areas of the enzyme where mutations on average led to increased activity are represented in green, while regions where mutations on average had a negative impact on activity are shown in red. The most frequent 6 positions among the top 1,000 mutants with highest normalized activity scores are indicated.

We finally deconvoluted the activity score from the expression score in order to identify regions of the enzyme susceptible to catalytic improvement. We first selected all single mutant variants with expression fitness values higher than the upper limit observed for nonsense mutants (Expression fitness = -0.69 ± 0.04; Fig. 2*F*). We then normalized the non-logarithmic activity fitness of 6,362 single mutants relative to their respective expression fitness values. These normalized activity scores exhibited reduced sensitivity to mutations’ impact on enzyme folding stability, as evidenced by the decrease in correlation with FOLDX algorithm predicted values, reduced from rho=0.59 (n=354, *p*<0.001, Fig. S4*D*) to rho=0.3 (n=354, *p*<0.001). Figure 5B shows the enzyme structured colored according to normalized activity. This normalization process resulted in 2,029 variants with normalized activity scores exceeding 1, indicating enhancements in catalytic activity independent of expression levels (Fig. S8*B, C*). We then ranked all variants based on their normalized activity scores and selected the top 1,000 variants from the entire dataset. We extracted the six most frequently represented amino acid positions within this subset and indicated them on the three-dimensional structure of the DAOx (Fig. 5*B*). Four out of the six identified positions were found to be closely situated to the substrate tunnel region, through which D-alanine approaches the enzyme and enters the active site (positions: 99, 216, 217, 218; Fig. 5*B*). In particular, S216, D217, and P218 are components of the loop positioned at the entrance of the tunnel and are in close proximity to the asparagine in position 99. These findings suggest that this region plays a role in controlling the accessibility of the substrate tunnel, through interactions with the substrate itself or by dynamically reshaping the enzyme’s structure to facilitate the entry of D-alanine into the reaction site. The other two positions identified, K109 and D110, are located on the opposite side of the dimer interface. Modifying these positions could potentially trigger conformational changes in the overall enzyme structure, which in turn may lead to allosteric effects on the enzyme’s catalytic function (Fig. 5*B*).

## Conclusions

EP-Seq is a deep mutational scanning workflow for studying enzyme folding stability and catalytic activity. Yeast surface display combined with peroxidase-mediated proximity labeling of single cells was able to link enzymatic activity to fluorescent phenotypes for large libraries in a pooled format, enabling cell sorting and high-throughput sequencing. We demonstrated our workflow by constructing a near comprehensive single substitution variant library of DAOx, and assaying it by EP-Seq for folding stability and catalytic activity. By jointly analyzing the expression and activity fitness datasets as a function of various biophysical and biochemical properties of the mutated residues at WT residues, we gained structural and biophysical insight into DAOx.

When considering the correlation of fitness score with various biophysical parameters of the mutated residues, for many parameters the two datasets exhibited high concordance. This shows how in order for a variant to be catalytically active, it must first be stably expressed and secreted to the cell wall. We therefore observed an expression level-dependency of the catalytic activity. However, our data further revealed certain properties of mutated sites which were differentially correlated with expression and activity fitness. These included distance of the mutated residue to the FAD cofactor and distance to the dimer interface. We found that WT residues in close proximity to the FAD cofactor and in the active site tended to destabilize the enzyme. These residues could be mutated to enhance folding stability, primarily through hydrophobic effects (Fig. 3*DI* and Fig. S8*E*). However, this increased stability was mostly achieved at the cost of catalytic activity (see Fig. 3D*IV* and Fig. S8*D*). All these observations comprehensively show how functional constraints have shaped the evolution of DAOx over time, and provide a direct and clear evidence of an activity-stability tradeoff. The distance of the mutated residues to the dimer interface was similarly decoupled between expression and activity fitness datasets. Prior literature^52,53^ indicates dimerization is necessary but not sufficient for catalytic activity. Mutations that disrupted the dimer interface therefore functioned similarly to those in the active site, where destabilizing the dimer interface through mutation could lead to higher overall expression, however this increased stability was achieved at the cost of catalytic activity. Our workflow is compatible with studying enzymes that can be directly or indirectly (via enzymatic cascade) linked to the production of peroxide. This includes enzymes with immediate therapeutic relevance such as Arginase^57^, and Asparaginase (Fig. S9), as well as biocatalysts with diagnostic or industrial applications, like Glucose Oxidase^28^.Due to its inherent scalability, our approach will find applications in generating training data for machine learning algorithms, which represents a major challenge for catalytic enzymes. In the future, EP-Seq can contribute to a better understanding of evolutionary processes in natural enzymes, help in identifying functional allosteric sites, and be used to evolve protein catalysts for industrial and biomedical applications.

## Acknowledgments

This work was supported by the University of Basel, ETH Zurich, the SNF-NCCR in Molecular Systems Engineering, and an SNSF Grant (200021_191962) to M.A.N. The authors thank the following individuals for helpful discussions: Joanan Lopez Morales, Jaime de Santaella, Olena Protsenko.

## Data availability statement

All the data required for replicating the study, DNA sequencing files and the raw data associated with each figure, have been deposited in the Zenodo repository and are accessible to the public through the DOI: 10.5281/zenodo.8388902. Additionally, the heatmaps and their associated raw data have been made available in color-blind accessible and interactive formats at the following location: https://nash-lab.github.io/DAOx-DMS/heatmaps/hm_expression_activity.html https://nash-lab.github.io/DAOx-DMS/heatmaps/hm_normalized_activity.html

## Code availability statement

All the custom computer codes and programs required to generate the data of this work are publicly available in the Zenodo repository and accessible through the DOI:10.5281/zenodo.8388902

