## Supplementary Information for "Understanding Activity-Stability Tradeoffs in Biocatalysts by Enzyme Proximity Sequencing"

### Methods

##### DAOx sequence cloning

The gene coding for *Rhodotorula gracilis* D-amino acid oxidase (DAOx) was acquired from Twist Bioscience codon optimized for expression in *Saccharomyces cerevisiae*. The sequence was cloned after restriction digestion with BamHI and XhoI into a pYDKan plasmid, a modified pYD1 yeast display vector where the original glycine-serine linker between Aga2 and the protein of interest was replaced with the protein linker GTPTPTPTPTGEF[58] and the β-lactamase gene replaced with a kanamycin resistance gene. After confirming the sequence through Sanger sequencing the plasmid pYDKan_RgDAOx was transformed through a lithium acetate transformation protocol[59] in the yeast strain EBY100 and the positive colonies selected on synthetic defined (SD) agar 2% (wt/vol) glucose plates lacking tryptophan (-Trp). We provide the sequence of the pYDKan_wtRgDAOx here**:**
https://zenodo.org/record/8388902

#### Expression and surface display of DAOx wild type

Positive transformants were cultivated in -Trp liquid medium with 2% (wt/vol) glucose for 24h at 30°C to an $OD_{600}$ ~8 with continuous shaking at 200 rpm. Expression and display of the Aga2_DAOx fusion protein were tested by transferring the cells to fresh -Trp liquid medium with 0.2% (wt/vol) glucose and 1.8% (wt/vol) galactose further supplemented with 100 mM citrate/phosphate buffer at pH7. In the optimization step of the protocol cells were transferred in an induction medium without buffer or with 100 mM citrate/phosphate buffer at pH range 3-7. Yeast cells were grown in induction media for 48 hours at 20°C before being pelleted, washed with PBS 0.1% (wt/vol) bovine serum albumin (BSA) and used for antibody labeling in order to detect the c-terminal 6xHistidine (6xHis) tag of the displayed fusion protein.

#### Yeast antibody staining to detect surface displayed DAOx

After induction of protein expression and display, yeast cells were washed twice with PBS 0.1% (wt/vol) BSA before being resuspended at a concentration of 2 million cells/100 µl in the same buffer with the addition of 1/500 dilution of anti-6x-His mouse monoclonal antibodies (Thermo Fisher, #MA1-21315). The samples were incubated with the primary antibody 30 min at room temperature before being washed twice with PBS 0.1% (wt/vol) BSA. After, the cells were resuspended at the concentration of 2 million cells/100 µl in PBS 0.1% (wt/vol) BSA with 1/500 dilution of a secondary goat anti-mouse antibody conjugated with Alexa Fluor™ 594 (Thermo Fisher #A-11005). The samples were incubated with the secondary antibody for 30 min at 4°C before being washed twice with PBS 0.1% (wt/vol) BSA and analyzed through flow cytometry. Surface expression of DAOx was determined through flow cytometry using a Attune NxT (Thermo Fisher Scientific) cytometer equipped with a 488 nm and a 561 nm laser. For flow cytometry, 10000 cells per sample were recorded and analyzed.

#### DAOx Amplex Red activity assay

Yeast cell populations positive for the display of DAOx were assayed through Amplex Red activity assay to detect D-amino acid oxidase activity. Half million yeast cells were mixed together with 35 mM D-alanine, 5.6 µM HRP, and 100 µM Amplex Red in PBS (pH 7.5). The fluorescence was read at 590 nm.

#### DAOx mutant library construction and barcoding

Mutagenesis of the DAOx wild type gene was performed through a plasmid based one-pot saturation mutagenesis protocol previously described by Wrenbeck and colleagues[60] . We transferred the entire DAOx expression cassette, inclusive of galactose promoter, Aga2- WT DAOx open reading frame (ORF) and Matα transcription terminator from pYDKan (5498 bp) to a smaller pUC19 plasmid (4265 bp). The new plasmid was further engineered through the insertion of two BbvCI restriction sites located upstream and downstream of the expression cassette. The original nicking mutagenesis protocol was integrated with an additional step: after the first incubation with Nt.BbvCI enzyme and exonucleases, the reaction mixture was further incubated with 10 units of Quick CIP phosphatase (New England Biolabs) at 37°C for 20 min, followed by incubation at 80°C for 20 minutes. This step promotes the removal of 5' phosphate from nicked and partially degraded strands of wild type DNA and prevents reformation of closed double strand wild type DNA plasmid in the following amplification and ligation steps. For the mutagenic DNA amplification step of the protocol we used nested NNK forward primers targeting codons 2 to 365 of the DAOx wild type sequence. Primers were designed using the script create_primers.py[61]. The length of each primer was adjusted within 25 and 51 nucleotides in order to obtain oligonucleotides with similar melting temperature in the range of 60-61°C. We ordered hand-mixed degenerate oligonucleotides in 96 well plate form from Integrated DNA technologies and mixed the primers in equimolar proportion at a final concentration of 10 µM. Finally, the one pot site saturation mutagenesis protocol was performed into two subsequent cycles. The pool of plasmids recovered after the first cycle was used as a template for the second cycle. Equimolar amounts of the resulting plasmids from the two rounds of mutagenesis were merged composing the final mutant library. Unique molecular identifier (15N) DNA barcodes were linked through standard PCR to each of the DAOx mutagenized inserts by using primers F1 and R1 (Table S3). Nucleotide barcodes were added to the 5' end of the amplicon, upstream of the galactose promoter. Amplified and barcoded expression cassettes with mutagenized DAOx coding sequences were cloned into a BbVCI/PmeI digested pYDKan vector using HiFi DNA Assembly (New England Biolabs). Assembled products were column purified (Zymo-Spin I, Zymoresearch) before being electroporated into XL1-Blue electroporation competent cells (Agilent, 200228). Positive colonies were selected on 15 cm LB agar plates containing kanamycin (50 µg/ml). The final barcoded library size was capped through single cell sorting to 200,000 cells. After sorting, cells were incubated for 20 hours and then used to extract plasmid DNA. 5 µg of the library plasmid pool was transformed into the final yeast *Saccharomyces cerevisiae* strain EBY100 through lithium acetate transformation[59]. Serial dilutions of the final transformation reactions were plated on -Trp agar plates with 2% (wt/vol) glucose. Quantification of the colonies forming units from the dilution plates indicated a final transformation yield between 6 and 10M colonies. After transformation, the yeast mutant library was grown for 24h at 30°C in -Trp liquid medium supplemented with 2% (wt/vol) glucose before being diluted to $OD_{600}$=1 into fresh -Trp medium supplemented with 2% (wt/vol) glucose and grown for further 16h at 30°C. Finally, aliquots of $10^8$ cells were prepared and resuspended in 25% (vol/vol) glycerol before being stored at -80°C. Sequences of pUC19_RgDAOx plasmid and mutagenic primers are provided here: https://zenodo.org/record/8388902

#### Pacific Biosciences long read sequencing and data analysis

We used PacBio long read sequencing to establish the link between the DAOx variants represented in our mutant library and the 15 nucleotides unique molecular identifiers (UMI). We sequenced the mutant library before transformation of the mutagenized and barcoded pool of plasmids into the final yeast strain EBY100. PacBio sequencing was performed on bacterial extracted plasmid DNA. This procedure eliminates the need of preparing the sequencing insert via PCR and thus the possibility of strand exchange events. Barcoded plasmids were purified from bacterial cells and the sequencing inserts prepared through restriction digestion with PmeI-NotI restriction enzymes. The final dsDNA pool was purified after agarose gel electrophoresis and used for SMRTbell ligation. PacBio sequencing run was performed on a Sequel IIe system with a 30h movie collection time using a SMRT cell 8M. Final PacBio circular consensus sequences (CCSs) were generated using the ccs program from Pacific Bioscience (https://github.com/PacificBiosciences/ccs) and filtered in order to retain DNA sequences with average Phred quality score higher than Q20. After this stage, the sequencing files from the two independent PacBio reactions were merged. The final merged sequencing file is available at https://zenodo.org/record/8388902

We established a computational workflow to map, extract and link each UMI represented in our library to the respective DAOx variant sequence. Through Minimap2[62] (version 2.19), using a reference sequence (DAOx_ref.fa), we mapped on each read the region corresponding to the UMI and the mutations falling in the Aga2-DAOx open reading frame.

**minimap2 --cs -ax map-hifi** DAOx_ref.fa pacbio.fastq **>** aln.sam

The output of Minimap2 alignment was processed further by using a C program called *ppba.c*. This program was specifically designed for this task and it makes use of a custom library file named *libsccodon.h*. Information regarding the nucleotide sequence of each barcode, barcodes sequencing quality per position, mutations falling in the Aga2-DAOx ORF and presence of indels were extracted and stored into a first version of the look-up table (pre_lut).

**./ppba** DAOx_ref.fa aln.sam **>** pre_lut.tsv

The resulting look up table file was processed to remove conflicting nucleotide barcodes through the jupyter notebook "Generate_Lut.ipynb". Identical UMIs associated with different DAOx variant sequences were compared. Among those, we retained in the final version of the look-up table UMIs whose link to a specific mutation was confirmed by more independent sequencing reads or UMIs with higher sequencing quality and linked to lower number of indels in the sequencing read. If none of these criteria could be used to classify the UMIs, conflicting barcodes were all removed from the final look-up table. UMIs linked to mutations registered out of the targeted region (codon 2-365 of DAOx) were kept in the lookup table and filtered out in a later stage of the analysis. The resulting look-up table file was further processed in order to finalize the table and remove all the nucleotide barcodes with length different than 15 nucleotides.

The C programs used for this analysis were compiled by using GCC (version 7.4.0) on a 32-bit processor. The scripts, sequencing files and the final look-up are available at https://zenodo.org/record/8388902

#### DAOx variant library expression and display

100M yeast cells transformed with the barcoded DAOx variant library were inoculated at starting OD600 =0.1 in 100 ml of -Trp liquid medium with 2% (wt/vol) glucose and grown for 24h at 30°C to an OD600 ~8 with continuous shaking at 200 rpm. Protein expression and display were induced by transferring the yeast cells at a starting OD600 of 0.4 to a 100 ml fresh -Trp liquid culture with 0.2% (wt/vol) glucose and 1.8% (wt/vol) galactose. The induction medium was further supplemented with 100 mM citrate-phosphate buffer at pH7. Cells were cultivated in the induction medium for 48 hours at 20°C shaking at 200 rpm, before being washed twice with 25 ml PBS 0.1% (wt/vol) BSA. 120M induced cells were distributed into 6 tubes at a concentration of 20M cells/ml. Detection of the displayed Aga2-DAOx protein construct was performed in PBS 0.1% (wt/vol) BSA incubating the cells with 1/500 dilution of the anti 6xHis-tag mouse monoclonal antibodies (Thermo Fisher, #MA1-21315) for 30 minutes at room temperature. After incubation with the primary antibody, cells were washed twice with ice cold PBS 0.1% (wt/vol) BSA buffer before being resuspended in the same buffer with 1/500 dilution of a secondary goat anti-mouse antibody conjugated with Alexa Fluor™ 594 (Thermo Fisher #A-11005). The reaction was then incubated at 4°C for 30 minutes. After the incubation the cells were washed twice and resuspended in ice-cold PBS 0.1% (wt/vol) BSA at a concentration of 20M cells/ml. In parallel, a negative control staining procedure was performed. In a single tube 20M induced yeast cells were incubated exclusively with the secondary goat anti-mouse antibody conjugated with Alexa Fluor™ 594 (Thermo Fisher #A-11005) at the same conditions described above. This sample was then used to detect eventual fluorescence signals caused by unspecific binding of the fluorescent secondary antibody to the yeast cells.

#### DAOx single cell tyramide/peroxidase proximity labeling assay

Catalytic activity of displayed DAOx wild type and variant enzymes was assayed through a single-cell tyramide/peroxidase proximity labeling method. After induction of protein expression and cell surface staining of the yeast population as described above, 25M yeast cells were mixed at a concentration of 500 cells/µl with 1/200 dilution of Alexa fluor™ 488 Tyramide Reagent (Thermo Fisher, B40953), 56.8 µM HRP (Sigma-Aldrich, 77332) and 130 mM D-alanine in 1xPBS and 0.75% (w/v) sodium alginate. The reaction was incubated for 20 minutes at 25°C. Afterwards, the cells were spun down for 3 min at 13000 g in a table top centrifuge. The cell pellet was washed twice with PBS 0.1% (wt/vol) BSA + 0.05% (vol/vol) Tween 20 and then used for flow cytometry and single cell sorting experiments.

#### Fluorescence activated cell sorting (FACS) of the DAOx yeast library

#### Expression level sorting.

Yeast cells stained for the expression and display of DAOx variants were sorted using a FACSMelody cell sorter equipped with 488 and 561 nm lasers and a 100 µm nozzle. Cells were sorted into pre-wet 5 ml FACS tubes containing 0.5 ml of 2X -Trp medium with 4% (wt/vol) glucose and 1% (wt/vol) BSA. Yeasts were first gated for single events and the population further divided into 4 sorting bins along the Alexa Fluor™ 594 fluorescence axis. The first bin (Bin1) was set in order to capture the 99% of non-fluorescent cells by using the negative control sample of the antibody staining procedure as reference. The remaining part of the cell population was divided into three further sorting bins capturing each an equivalent fraction of the yeast population with increasing fluorescence intensity (Bins 2,3,4). The sorting procedure was repeated two independent times sorting each time more than 10M single yeast cells (Table S2).

#### Activity level sorting.

EBY100 yeast cells stained for the expression of the DAOx enzyme variants and assayed through single cell tyramide activity assay were sorted using a FACSMelody cell sorter equipped with 488 and 561 nm lasers and a 100 µm nozzle. As above, cells were sorted into pre-wet 5 ml FACS tubes containing 0.5 ml of 2X -Trp medium with 4% (wt/vol) glucose and 1% (wt/vol) BSA. Yeast cells were first gated for singleton events and then the population divided into four bins based on the level of green fluorescent signal. Bin 1 was designed in order to include 99% of the population of cells negative to display or displaying inactive DAOx variants, using as reference the fluorescence level of the non-displaying yeast populations. The remaining part of the population of cells was equally divided into three yeast sub-populations with increasing fluorescent signal. We performed the sorting of two independently assayed yeast populations sorting each time more than 8 M total yeast cells (Table S2).

After each cycle of sorting both for expression and activity, yeast cells part of the same gated population were merged into 50 ml falcon tubes and pelleted 10 min at 4000 g in a table top centrifuge. Afterwards, the supernatant was discarded and the cell pellet resuspended in 10 ml -Trp medium with 2% glucose supplemented with 100 µg/ml Penstrep. After sorting, all the cell populations were grown for 48h at 30°C shaking at 200 rpm before being sampled into aliquots of 50M cells each and stored at -80°C in 25% (v/v) glycerol.

#### DNA prep for Illumina sequencing

50M yeast cells per sorted population were used as starting material for the preparation of Illumina sequencing inserts. Cells were first collected from -80°C and incubated 5 minutes at RT before being spun down 1 min at 13,000 g in a table top centrifuge. The supernatant was discarded and the cell pellet resuspended in 250 µl of miniprep resuspension solution (GeneJET Plasmid Miniprep Kit, Thermo Fisher) with the addition of 4 µl of Zymolyase (5U/µl). The reaction was incubated for 2h at 37°C shaking at 900 rpm. After this incubation step the samples were processed following a typical plasmid miniprep kit protocol (GeneJET Plasmid Miniprep Kit, Thermo Fisher). Finally, plasmid DNA extracted from yeast cells was eluted in 15 µl of nuclease free water. The region of the plasmids containing the 15N UMI was amplified through a standard PCR reaction using the NEBNext Ultra II Q5 Master Mix (New England Biolabs). 25 µl of the master mix were mixed with 5 µl of each primer (1 µM) and 15 µl of template DNA. Primers were designed accordingly in order to target the region of interest and be compatible with the Nextera indexing library preparation. An equimolar mixture of four staggered primers was used in each sample preparation as forward and reverse primer in order to provide in both ends of the resulting amplicon different starting nucleotides for the Illumina reads (F3-6 and R3-6 primers in Table S3). PCR was performed with the following program: 1 cycle at 98°C for 30 sec, 18 cycles at 98°C for 10 sec, 72°C for 30 sec, 72°C for 2 min and a final elongation step of 2' at 72°C before storing the reaction at 4°C. Resulting DNA amplicons were then visualized through DNA electrophoresis on a 1% (wt/vol) agarose gel in and purified from the gel using a standard DNA purification kit. The concentration of DNA per sample was measured and DNA was purified once more through the DNA Clean and Concentrator-5 kit (Zymoresearch). After addition of unique Nextera indexing sequences the samples were pooled and sequenced through Novaseq 6000 Illumina sequencing. Numbers of Illumina reads per sample are provided in Table S2.

Demultiplexed reads were then processed through a computational pipeline in order to extract the sequences of UMIs and align them to the information contained in the look-up table.
Sequences corresponding to the UMI were first mapped through BBMap alignment algorithm [63] by using a reference sequence file (ill_ref.fa)

```
bbmap.sh in= *.fastq ref=ill_ref.fa out= *.sam
```

BBMap *.sam outputs were further processed through the C script *pib.c* (process illumina barcodes). Through this step we extracted the sequences and read quality of each UMI mapped by BBMap, saving the information in a .fastq format file.

```
./pib *.sam > *.fq
```

Each UMI mapped and registered was then aligned to the information stored in the look-up table through the C script *rib.c* (read illumina barcodes). The following tags: **0** - not found in the look-up table, **1** - found in the look-up table, **2** - read quality < Q20, **3** - UMI size different than 15N, were associated with each of the barcodes read through Illumina sequencing. We applied a quality filter of Q20, therefore tagging with tag 2 all the UMI associated with a read quality lower than Q20.

```
./rib -t lut_m.tsv -q 20  *.fq  > *.tsv
```

Finally, UMI sequences with tag 1 (found in the look-up table), were extracted, sorted alphabetically and grouped by identity.

```
grep ^1 *.tsv|sort|uniq -c|sed -E 's/^ *//; s/ /\t/'> t1sct_*.tsv
```

All the Illumina reads files were processed through the reported computational workflow. Finally all the information about the UMI sequences and count found in each bin (t1sct_Bin#.tsv) were merged in a unique file using the jupyter notebook "ill_tag1_bins".
The C programs used for this analysis were compiled by using GCC (version 7.4.0) on a 32-bit processor.
The scripts and sequencing files are available at https://zenodo.org/record/8388902

#### Expression and activity fitness scores calculation

The number of sequencing reads linked to each variant enzyme ($r_v$) was converted into number of sorted cells of the same variant ($c_v$) per sorted bin using the following equation 1:

$$\frac{r_v}{r_{tot}} = \frac{c_v}{c_{tot}} \qquad (1)$$

where $r_{tot}$ is the total number of illumina reads of the bin and $c_{tot}$ is the total number of cells sorted in the same bin. Then, the final expression and activity scores per variant were computed as the expected value of fluorescent intensity of the variant across all the four bins of the experiment. We calculated first a weighted mean ($\beta$) of the cell numbers ($c_v$) where the median fluorescence of the yeast population in each bin ($\omega$) was used as the weighting factor:

$$\beta = \frac{\sum_{i=1}^{bin} \omega_i \cdot c_{vi}}{\sum_{i=1}^{bin} c_{vi}} \quad \textbf{(2)}$$

The expression and activity fitness score ($F$) per variant were finally calculated as follow:

$$\mathrm{F} = \log_2\left(\frac{\beta_v}{\beta_{wt}}\right) \quad \textbf{(3)}$$

where $\beta_v$ is the weighted mean expression score of the variant enzyme and $\beta_{wt}$ is the score of wild type DAOx. To calculate the final consensus fitness score ($F_{fin}$) for each variant enzyme, we used a weighted mean of the single fitness scores recorded in each replicate ($F_v$) of both the expression and activity assays. The weighting factor was the number of cells associated with the measured fitness in each experiment ($c_v$) (Equation 4).

$$F_{fin} = \frac{\sum_{j=1}^{rep} \mathrm{F}_{vj} \cdot c_{vj}}{\sum_{j=1}^{rep} c_{vj}} \quad (4)$$

#### FoldX ΔΔG calculation

In order to predict the effect of missense mutations on the stability of the DAOx monomeric structure we used the FOLDX algorithm[48] (version 5.0). We first processed the DAOx crystal structure (PDB:1C0P) through the FOLDX "RepairPDB" function in order to identify and repair residues of the structure with bad torsion angles or energy clashes. The repaired structure was then processed through the FOLDX tool "PositionScan" that mutates the residue of the structure (positions 2-361) to the 20 natural amino acids providing an estimation of the change in energy between the folded and unfolded state of the wild type protein and comparing it to the change in folding energy upon amino acid mutation. The final score ΔΔG was then calculated through the following equation:

$$\Delta\Delta G = \Delta G\text{Mut} - \Delta G\text{Wt} \quad \textbf{(5)}$$

Positive values of predicted ΔΔG scores indicate a negative effect on the stability of the structure. Negative values of ΔΔG predictions indicate positive effect of mutations on the overall stability of the structure. A mean ΔΔG score per position of the protein was calculated and the final values (n=360) correlated to experimental expression and activity fitness scores. Input-output and setting files used for the ΔΔG prediction are provided at https://zenodo.org/record/8388902

#### DAOx structural properties calculation and extraction

Solvent accessible surface area (SASA) was used as a measure of residue solvent exposure. SASA scores were calculated per residue of the DAOx wild type monomeric protein (PDB: 1C0P) through the PyMol tool "get_sasa_relative" (https://pymolwiki.org/index.php/Get_sasa_relative). Scores ranging between 0 (minimum solvent exposure) and 100 (maximum solvent exposure) were extracted and analyzed in relation to the activity and expression fitness scores.

B-factor scores as a scale of thermal induced dynamic disorder and flexibility of the structure were extracted per position of DAOx monomeric structure (PDB file 1C0P.pdb).
Residues involved in the monomer-monomer contact at the dimer interface of DAOx dimeric structure (PDB:1C0P) were selected through the pyMol tool "InterfaceResidues" (https://pymolwiki.org/index.php/InterfaceResidues ).
Coordinates of the alpha carbons (Cα) of interface residues were extracted from the file 1C0P.pdb and the distance between each residue of the structure to the closest interface Cα was calculated. Spatial coordinates of all the atoms of the flavin adenine dinucleotide (FAD) cofactor were extracted from the 3D structure of DAOx wild type (PDB:1C0P). We calculated the distance of each Cα of DAOx monomeric protein to each of the FAD cofactors atoms. A final mean value distance per residue to the FAD was calculated by computing the arithmetic mean of all the distances of the same residue to the FAD atoms.

#### Single mutant variants expression and activity level

From the 6387 single mutant variants of DAOx analyzed through our DMS workflow both for expression and activity, we randomly selected 12 single mutant DAOx variants: S48C, L153R, A187K, A187E, G199E, Q200W, S215A, T237M, S268Y, P292E, L310P, G315P. Genes coding for each of the selected DAOx variants and codon optimized for the expression in *S.cerevisiae* were synthesized and cloned into a recipient pYDKan plasmid in frame with Aga2p coding sequence on a BioXP 3250 synthetic biology workstation (Codex DNA). All the sequences were confirmed through Sanger sequencing and the final plasmids transformed into the yeast strain EBY100 through lithium acetate transformation protocol. Positive colonies were selected on synthetic defined (SD) agar 2% (wt/vol) glucose plates lacking tryptophan (-Trp). Positive colonies were cultivated in -Trp liquid medium with 2% (wt/vol) Glucose for 24h at 30°C to an OD600 ~8 with continuous shaking at 200 rpm. Expression and display of the Aga2-DAOx wild type and mutant constructs were induced by transferring the cells at OD600=0.4 to fresh -Trp liquid medium with 0.2% (wt/vol) glucose and 1.8% (wt/vol) galactose and supplemented with 100 mM citrate/phosphate buffer at pH7. The cells were grown in induction medium for 48 hours at 20°C before being pelleted, washed with PBS 0.1% (wt/vol) BSA and used for antibody labeling in order to detect the c-terminal 6xHistidine (6xHis) tag of the displayed fusion protein.

#### Single mutant variants expression level.

Yeast cells stained for the expression of DAOx wild type and mutant variants were analyzed through flow cytometry. Yeasts were first gated for single events and the resulting population of cells was further divided into 4 gates along the Alexa Fluor™ 594 fluorescence axis. Gate 1, set between fluorescence values 0-2500, included the 99% of the non-stained yeast population used as negative control. Other 3 gates were set respectively at 2500-25000, 25000-250000, 250000-2500000 fluorescence arbitrary units in order to cover the entire range of fluorescence. 10'000 single cells from each yeast population were analyzed. We recorded the median value of the population in each of the four gates and the % of yeast cells represented in each gate. Each population was assayed for expression three independent times. We calculated a weighted mean (β) of the distribution of cells (%c) where the median fluorescence of the yeast population in each gate (ω) was used as the weighting factor (equation 6).

$$\beta = \frac{\sum_{i=1}^{gate} \omega_i \cdot \%c_i}{\sum_{i=1}^{gate} \%c_i} \qquad (6)$$

Finally, the single clone expression score ($F_{sing.\ clone}$) was calculated by dividing the weighted mean expression of each DAOx variant tested ($\beta_v$) by the weighted mean of the wild type DAOx ($\beta_{wt}$) assayed through the same procedure (equation 7)

$$F_{sing.\ clone} = \frac{\beta_v}{\beta_{wt}} \qquad \mathbf{(7)}$$

#### Single mutant variants activity level.

To measure the activity of DAOx wild type and mutant variants, yeast cell populations expressing these variants were assayed using the Amplex Red method with D-alanine as the substrate. A mixture of half a million yeast cells, 35 mM D-alanine, 5.6 µM HRP, and 100 µM Amplex Red in PBS (pH 7.5) was prepared. The fluorescence signal was then measured at 590 nm every 60 seconds for a total of 30 minutes. The linear range of the reaction for all the variant enzymes was determined between 0 and 10 minutes of incubation, and the slope of each reaction was calculated. The single clone activity score for each variant was obtained by dividing the slope of the reaction for the variant by the slope of the reaction for the wild type enzyme. The final scores were calculated as the average of six independent measurements per cell population.

#### Supplementary tables

##### Table S1. Look-up table composition.

Nucleotide and amino acid DAOx variants (codons 2-365) registered in the mutant library and associated with unique molecular identifiers (UMI).

| | **# Variants** | **# UMI** (tot: 173,002) |
|---|---|---|
| 0 nucleotide mutations (WT) | - | 54,158 |
| 1 nucleotide mutations | 6,503 missense | 59,743 |
| | 336 nonsense | 2,689 |
| | 308 synonymous | 4,692 |
| 2 nucleotides mutations | 31,943 missense | 33,812 |
| | 2,814 nonsense | 2,983 |
| | 171 synonymous | 178 |
| 3-5 nucleotides mutations | - | 14,747 |

#### Table S2. Number of sorted cells and Illumina reads.

Information about the number of cells sorted per bin in each of the expression level and activity level screening experiments. Number of Illumina reads per sample and median fluorescence values per sorting gate are also provided.

| Experiment | Bin | Median fluorescence | # sorted cells | # Illumina reads |
|---|---|---|---|---|
| Expression level Replicate 1 | # 1 | 146 | 6,243,697 | 42,024,431 |
| | # 2 | 3,424 | 1,926,870 | 39,848,257 |
| | # 3 | 19,202 | 1,757,354 | 41,267,548 |
| | # 4 | 48,439 | 1,704,305 | 41,474,350 |
| Expression level Replicate 2 | # 1 | 145 | 6,662,430 | 41,086,125 |
| | # 2 | 1,938 | 2,033,831 | 32,102,375 |
| | # 3 | 14,289 | 1,739,456 | 40,933,088 |
| | # 4 | 40,094 | 1,554,250 | 42,280,563 |
| Activity level Replicate 1 | # 1 | 825 | 5,526,601 | 41,466,216 |
| | # 2 | 4,873 | 814,188 | 38,937,993 |
| | # 3 | 15,295 | 1,001,782 | 36,382,700 |
| | # 4 | 40,787 | 908,535 | 38,617,574 |
| Activity level Replicate 2 | # 1 | 812 | 6,454,983 | 38,514,974 |
| | # 2 | 4,897 | 837,272 | 41,667,242 |
| | # 3 | 14,973 | 1,027,386 | 42,046,908 |
| | # 4 | 42,012 | 933,111 | 40,046,573 |

**Table S3. DNA primers used in this work.**

Sequence of NNK primers used to generate the mutant RgDAOx library are provided at: https://zenodo.org/record/8388902

| Primer name | 5'-3' sequence |
|---|---|
| F1 | GCGCGGCCTTTTGCCCTGCAGGCCNNNNNNNNNNNNNNNAGGGAACAAAAGCTGGCTAGTACGG |
| R1 | CGATTTTGTTACATCTACACTGTTGTTATCAGATCAGCGGGTTTAAAC |
| F3 | TCGTCGGCAGCGTCAGATGTGTATAAGAGACAGGGCCTTTTGCCCTGCAGGC |
| F4 | TCGTCGGCAGCGTCAGATGTGTATAAGAGACAGTGGCCTTTTGCCCTGCAGGC |
| F5 | TCGTCGGCAGCGTCAGATGTGTATAAGAGACAGATGGCCTTTTGCCCTGCAGGC |
| F6 | TCGTCGGCAGCGTCAGATGTGTATAAGAGACAGCATGGCCTTTTGCCCTGCAGGC |
| R3 | GTCTCGTGGGCTCGGAGATGTGTATAAGAGACAGGGAGGAGAGTCTTCCTTCGGAGGG |
| R4 | GTCTCGTGGGCTCGGAGATGTGTATAAGAGACAGTGGAGGAGAGTCTTCCTTCGGAGGG |
| R5 | GTCTCGTGGGCTCGGAGATGTGTATAAGAGACAGATGGAGGAGAGTCTTCCTTCGGAGGG |
| R6 | GTCTCGTGGGCTCGGAGATGTGTATAAGAGACAGGATGGAGGAGAGTCTTCCTTCGGAGGG |

#### Table S4. Effect of mutant amino acid on expression and activity scores.

AR= aromatic; CN= charge negative; CP= charge positive; HP= hydrophobic; PU= polar uncharged; SC= special case

| Mutant amino acid | Amino acid class | Expression fitness | Expression fitness (% change from average) | Activity fitness | Activity fitness (% change from average) | Count |
|---|---|---|---|---|---|---|
| Y | AR | -0.200 | -11.754 | -0.374 | -11.304 | 310 |
| F | AR | -0.197 | -9.711 | -0.354 | -5.393 | 314 |
| W | AR | -0.232 | -29.113 | -0.393 | -16.806 | 332 |
| D | CN | -0.180 | -0.633 | -0.391 | -16.353 | 317 |
| E | CN | -0.159 | 11.321 | -0.373 | -10.905 | 308 |
| R | CP | -0.241 | -34.273 | -0.407 | -20.999 | 337 |
| H | CP | -0.180 | -0.273 | -0.329 | 2.016 | 300 |
| K | CP | -0.218 | -21.621 | -0.391 | -16.460 | 328 |
| V | HP | -0.143 | 20.226 | -0.280 | 16.788 | 328 |
| M | HP | -0.133 | 25.550 | -0.264 | 21.533 | 318 |
| A | HP | -0.148 | 17.482 | -0.273 | 18.700 | 317 |
| G | HP | -0.190 | -5.725 | -0.345 | -2.689 | 321 |
| I | HP | -0.149 | 16.780 | -0.269 | 19.823 | 298 |
| L | HP | -0.172 | 4.024 | -0.290 | 13.631 | 325 |
| N | PU | -0.164 | 8.523 | -0.318 | 5.399 | 313 |
| Q | PU | -0.164 | 8.757 | -0.322 | 4.305 | 306 |
| S | PU | -0.148 | 17.291 | -0.275 | 18.238 | 335 |
| T | PU | -0.139 | 22.753 | -0.283 | 15.708 | 333 |
| P | SC | -0.281 | -56.884 | -0.527 | -56.839 | 328 |
| C | SC | -0.148 | 17.284 | -0.263 | 21.605 | 331 |
| | AR | -0.210 | -16.859 | -0.374 | -11.168 | 956 |
| | CN | -0.170 | 5.344 | -0.382 | -13.629 | 625 |
| | CP | -0.213 | -18.722 | -0.376 | -11.814 | 965 |
| | HP | -0.156 | 13.056 | -0.287 | 14.631 | 1907 |
| | PU | -0.154 | 14.331 | -0.299 | 10.912 | 1287 |
| | SC | -0.215 | -19.800 | -0.395 | -17.617 | 659 |

#### Supplementary figures

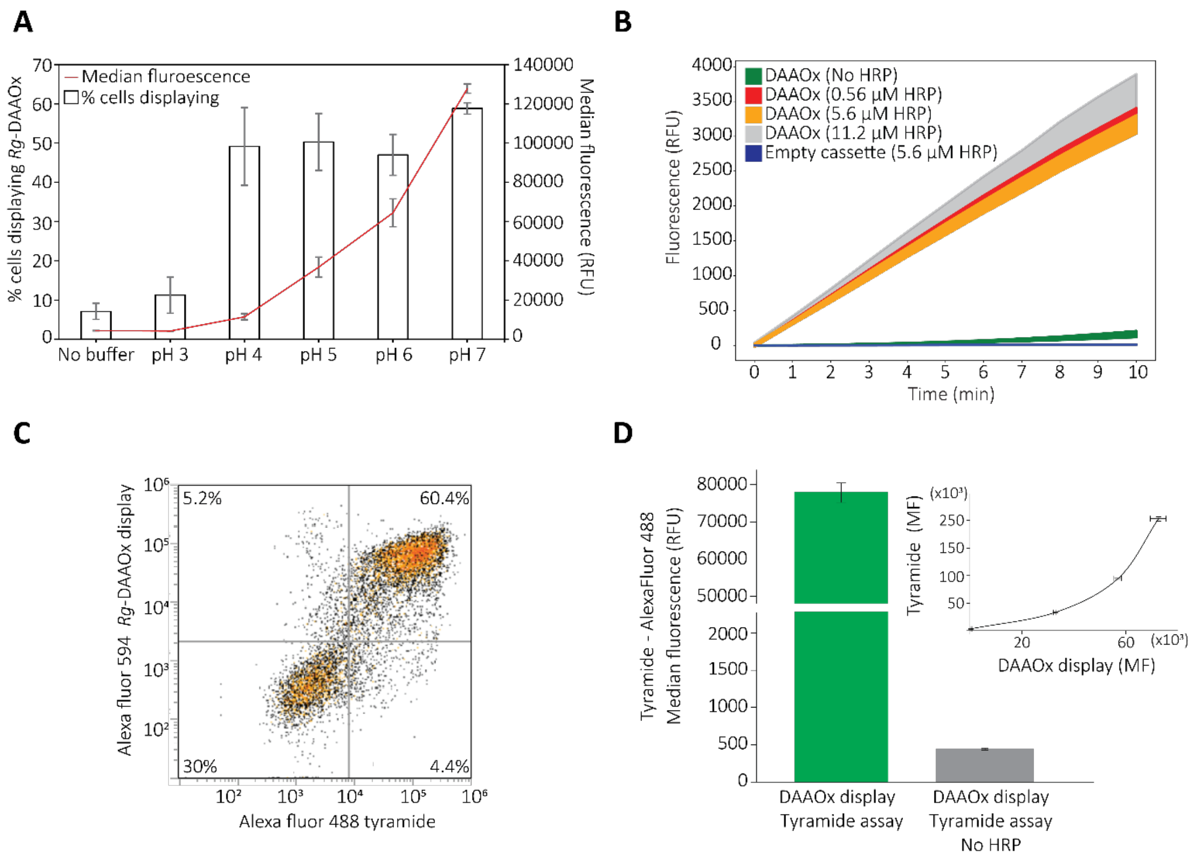

**Figure S1. Yeast surface display of D-amino acid oxidase from *Rhodotorula gracilis*.**
A) DAOx surface display levels after induction of protein expression at 20°C for 48h in growth media at different pHs. Percentage (%) of yeast population displaying DAOx wild type and median fluorescence relative fluorescent units (RFU) of the displaying population are reported. B) Population based DAOx activity assay. Half million cells from a DAOx displaying population were used for an Amplex Red activity assay to detect the activity of the displayed enzyme. A reaction using cells not displaying DAOx and a reaction using DAOx displaying cells but not HRP in the mixture were used as negative controls. The reaction was tested at increasing concentration of HRP in order to verify that HRP concentration was not limiting the rate of the reaction. C) Single cell tyramide activity assay. A population of cells positive for the display of the DAOx wild type enzyme was assayed for the activity of the enzyme through a single cells tyramide assay. The same assay was then used for the screening of the DAOx variant library. D) Tyramide-AlexaFluor 488 fluorescence of a cell population displaying DAOx incubated with the complete tyramide assay reaction mixture and with the reaction mixture lacking HRP. The inset shows how the median fluorescence (MF) linked to the surface display of DAOx correlates with the fluorescence gained by the cells assayed through tyramide assay.

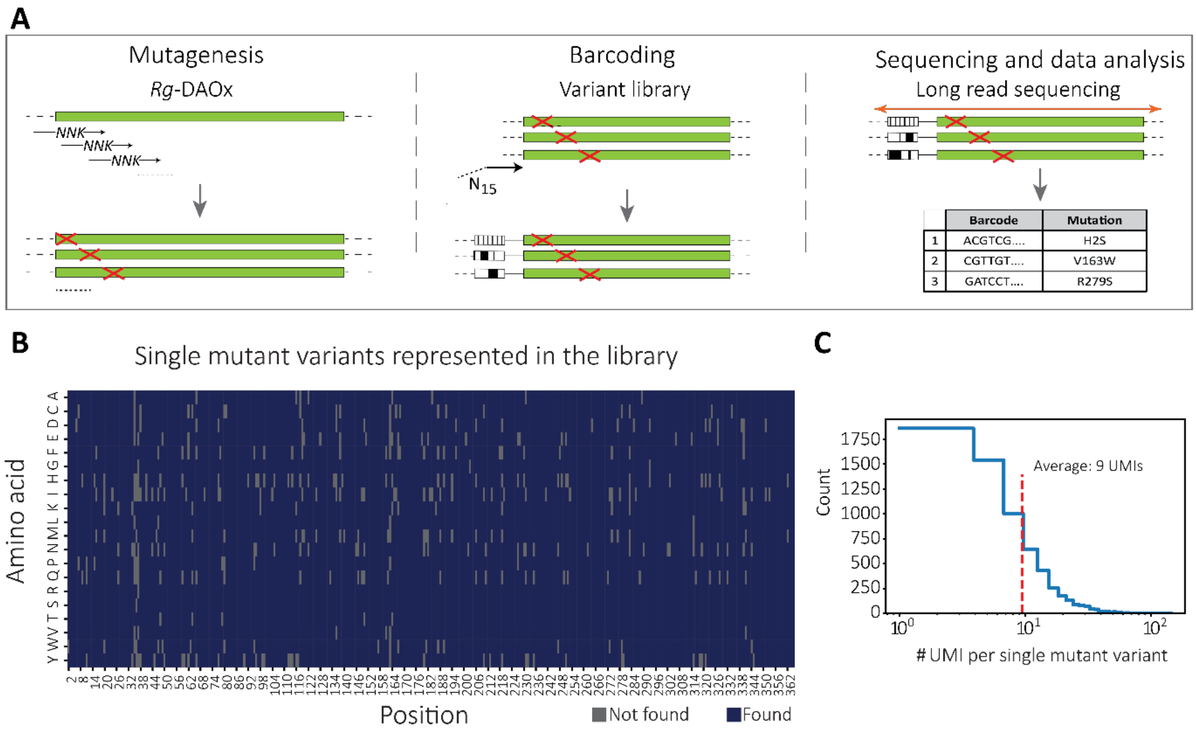

**Figure S2. Mutant library construction and sequencing.**
(A) A variant library of the enzyme DAOx was constructed by means of one pot site saturation mutagenesis[60]. Variants represented in the library were linked to unique molecular identifiers (UMI) by PCR. PacBio long read sequencing was used to reveal the pairs variant-barcode and the information stored in a look up table. (B) 6530 of 6916 possible single amino acid missense mutants are represented in the library (94.4% of total). (C) Number of UMIs associated with single mutant missense variants represented in our library.

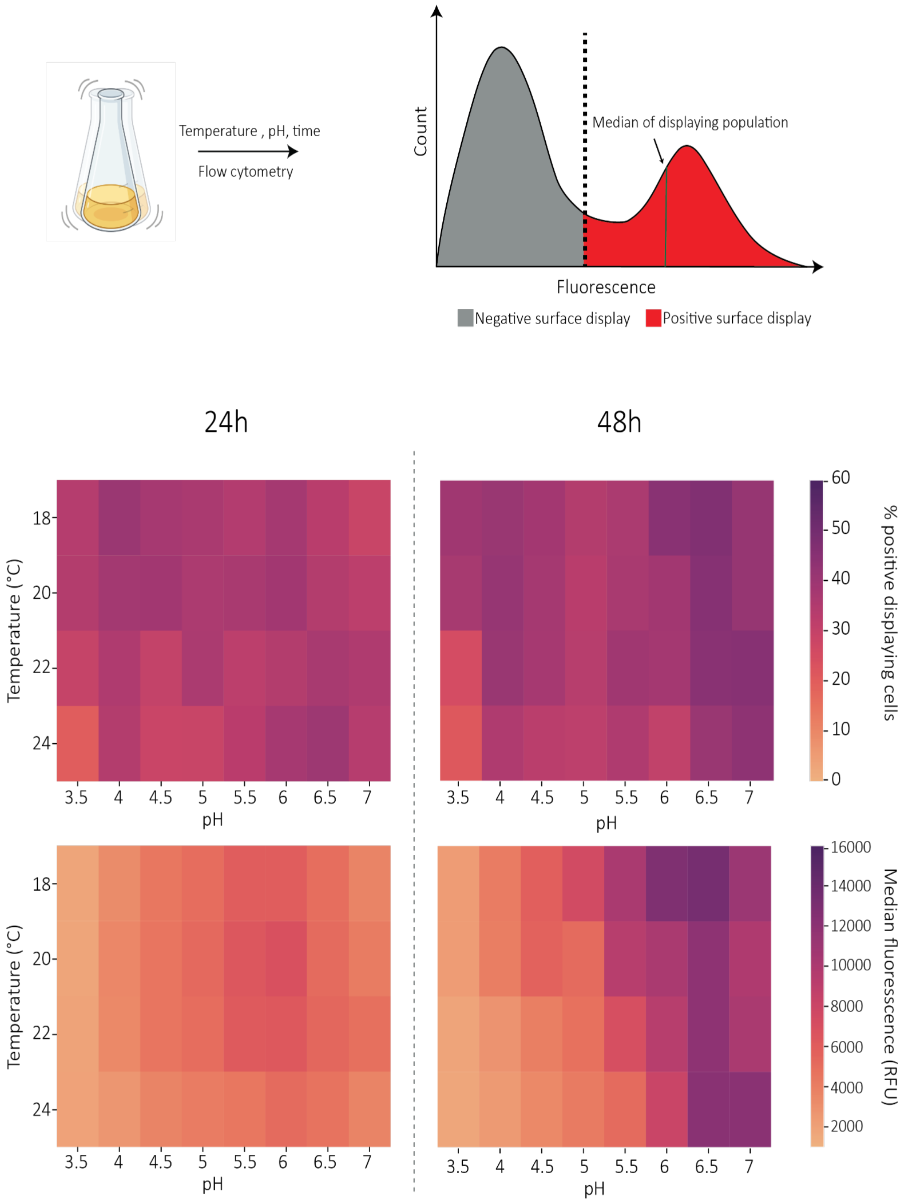

**Figure S3. Yeast surface display optimization of DAOx mutant library.**
The expression and display level of the variant enzymes were tested at 24 and 48h and at increasing temperatures (from 18 to 24°C) and pHs (3.5 to 7). The display of DAOx on the surface of yeast cells was detected through fluorescent antibody staining by targeting the C-terminal histidine tag of the Aga2-DAOx fusion construct. The percentage and median fluorescence of the positive displaying population were measured and are represented as heatmaps. The "peak" condition for both the evaluation criteria appears at 48h of protein induction time, pH range 5 to 7 and temperature range between 18 and 20 °C.

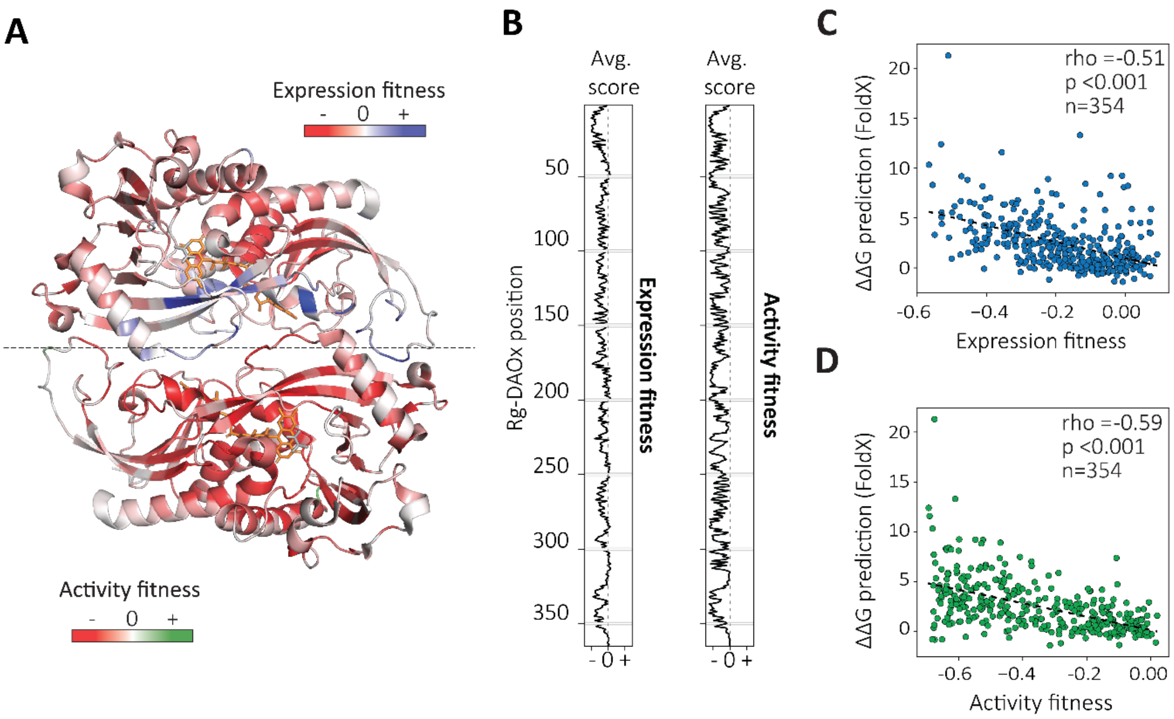

**Figure S4. Mean fitness expression and activity scores and their correlation with predicted ΔΔG.**

A) Mean expression (top) and activity (bottom) fitness scores per position of the DAOx mapped on the 3D structure of the enzyme (PDB 1C0P). B) Average expression (left) and activity (right) fitness scores along the sequence of DAOx. C) Spearman's rank correlation between predicted ΔΔG values and experimental expression scores per position of the protein. D) Spearman's rank correlation between predicted ΔΔG values and experimental activity scores per position of the protein.

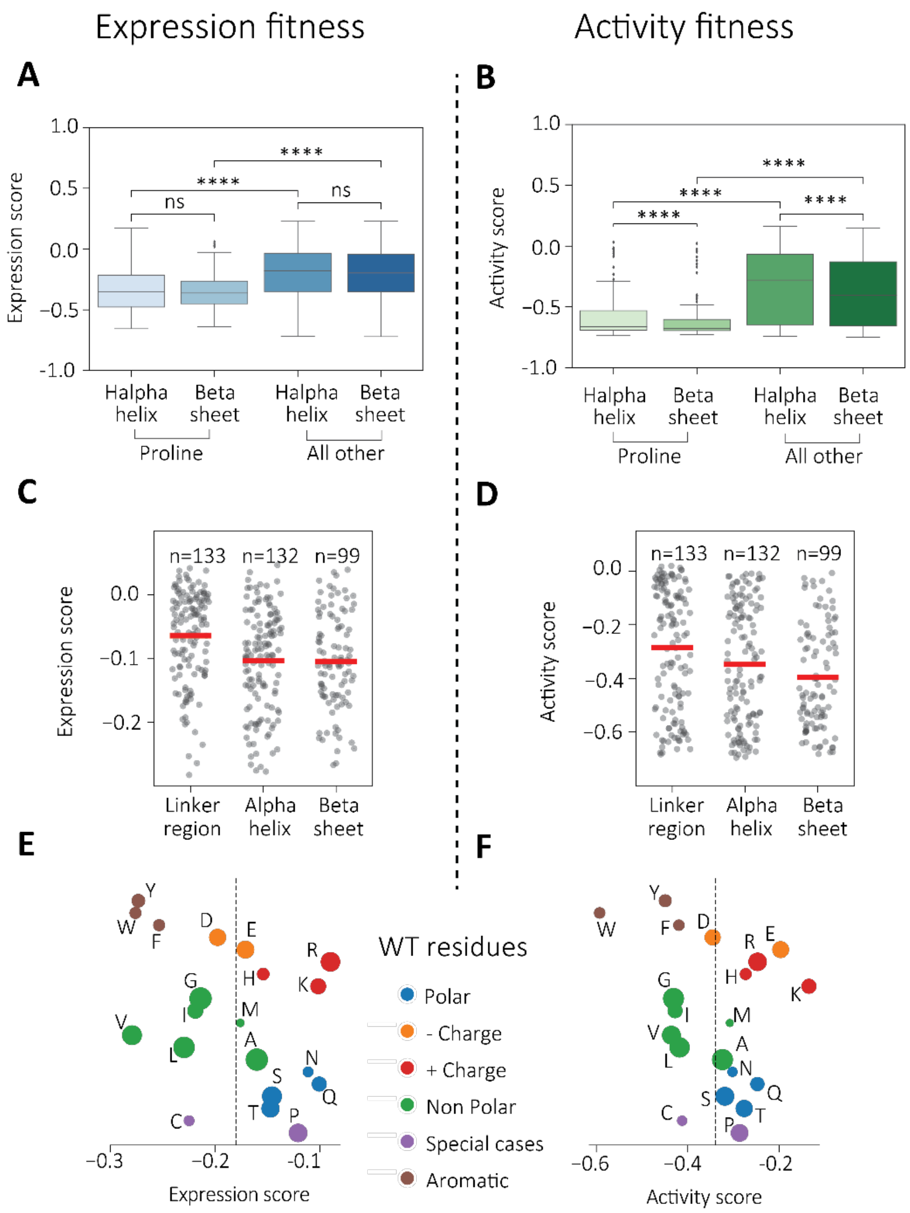

**Figure S5. Analysis of mutational effects through biochemical properties of amino acids and structural elements of DAOx enzyme.**

A) Impact of proline insertion on the expression of DAOx. B) Impact of proline insertion on the enzymatic activity of DAOx. C) Effect of single amino acid mutations on the expression of DAOx in relation to its secondary structure elements. D) Effect of single amino acid mutations on the enzymatic activity of DAOx in relation to its secondary structure elements. E) Impact of the substitution of wild type amino acids on the expression of DAOx. F) Impact of the substitution of wild type amino acids on the activity of DAOx. Size of the dots represents a relative measure of the amino acid abundance in the wild type sequence of DAOx. The reference fitness score (dashed line) was established as the average fitness values derived from grouping the dataset by the nature of substituted wild-type amino acids, resulting in -0.182 for expression and -0.342 for activity.

A

| Property | Expression score dataset Pearson *r* , p-value (*p*) | | Activity score dataset Pearson *r* , p-value (*p*) | |
|---|---|---|---|---|
| WT Bulkiness | **-0.10** | *p* <0.001 | **-0.07** | *p* <0.001 |
| Mut Bulkiness | **-0.03** | *p* =0.03 | **-0.01** | *p* =0.43 |
| Δ Bulkiness | **-0.06** | *p* <0.001 | **-0.05** | *p* <0.001 |
| WT Polarity | **0.19** | *p* <0.001 | **0.25** | *p* <0.001 |
| Mut Polarity | **-0.02** | *p* =0.11 | **-0.08** | *p* <0.001 |
| Δ Polarity | **0.14** | *p* <0.001 | **0.22** | *p* <0.001 |
| WT Hydrophob. | **-0.22** | *p* <0.001 | **-0.26** | *p* <0.001 |
| Mut Hydrophob. | **0.06** | *p* <0.001 | **0.09** | *p* <0.001 |
| Δ Hydrophob. | **-0.20** | *p* <0.001 | **-0.25** | *p* <0.001 |

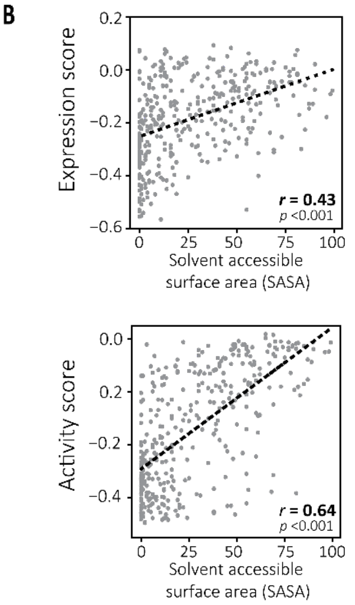

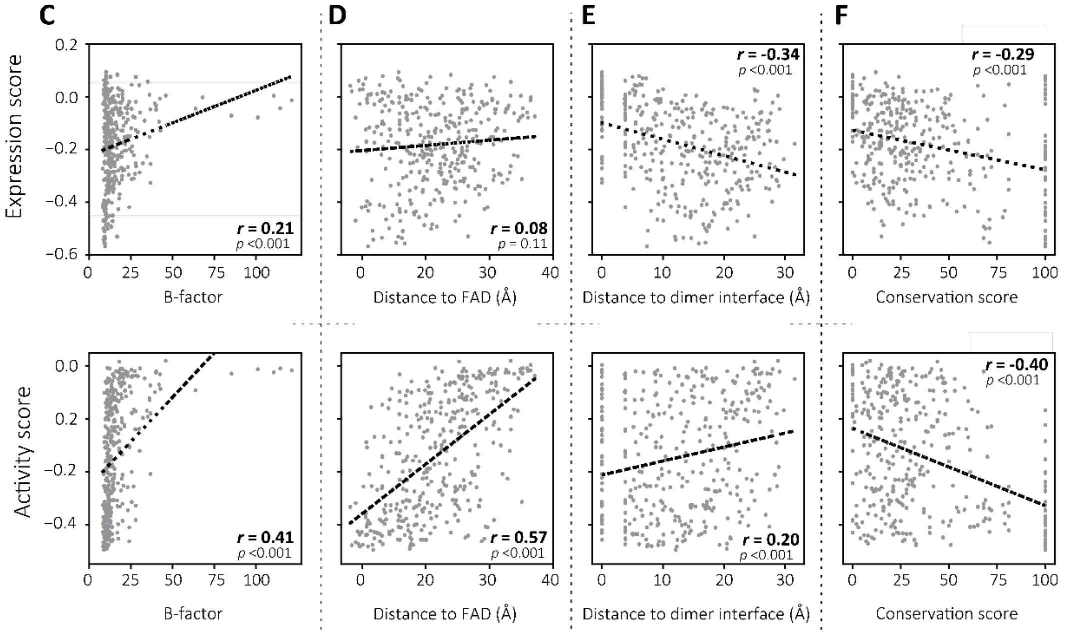

#### Figure S6. Linear correlation of the expression and activity DAOx datasets with biochemical, structural and functional properties of the enzyme

A) Pearson correlation coefficient (*r*) of activity and expression datasets (n=6399) with chemical physical properties of wild type and mutant amino acids. B) Linear regression of the expression (top) and activity (bottom) datasets with solvent accessible surface area (SASA) score calculated per position of the protein (n=360). C) Linear regression of the expression (top) and activity (bottom) fitness scores with temperature factor (B-factor) extracted from the crystal structure data of DAOx (n=360). D) Linear regression of the expression (top) and activity (bottom) scores with the distance of the mutation site from the FAD cofactor (n=360). E) Linear regression of the expression (top) and activity (bottom) fitness scores with the distance of the mutation site to the dimer interface. All the structural features were extracted from the 3D structure of DAOx (n=360) (PDB: 1C0P). F) Linear regression of the experimental expression and activity datasets with natural evolution conservation score calculated by aligning the wild type sequence of DAOx with 5 evolutionary related DAOx protein sequences (n=360) (see Fig. S7).

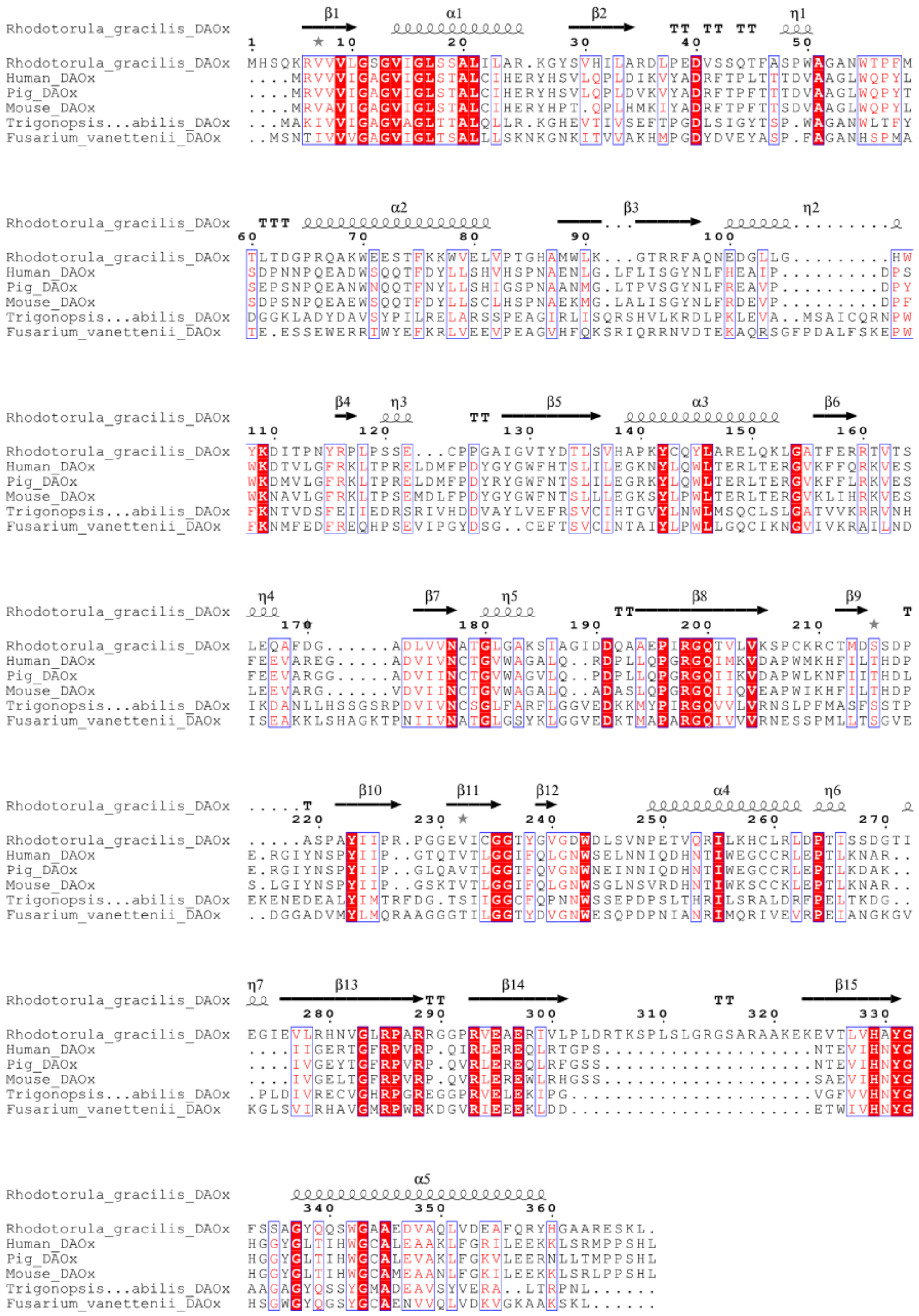

#### Figure S7. DAOx sequence alignment

Alignment of the protein sequence of *Rhodotorula gracilis* DAOx studied in this work, to evolutionary related D-amino acid oxidase sequences from human, pig, mouse, *Trigonopsis variabilis* and *Fusarium vanettenii* [64]. ClustalX software was used to compute the alignment and extract a conservation score per position of the DAOx protein from *Rhodotorula gracilis.*

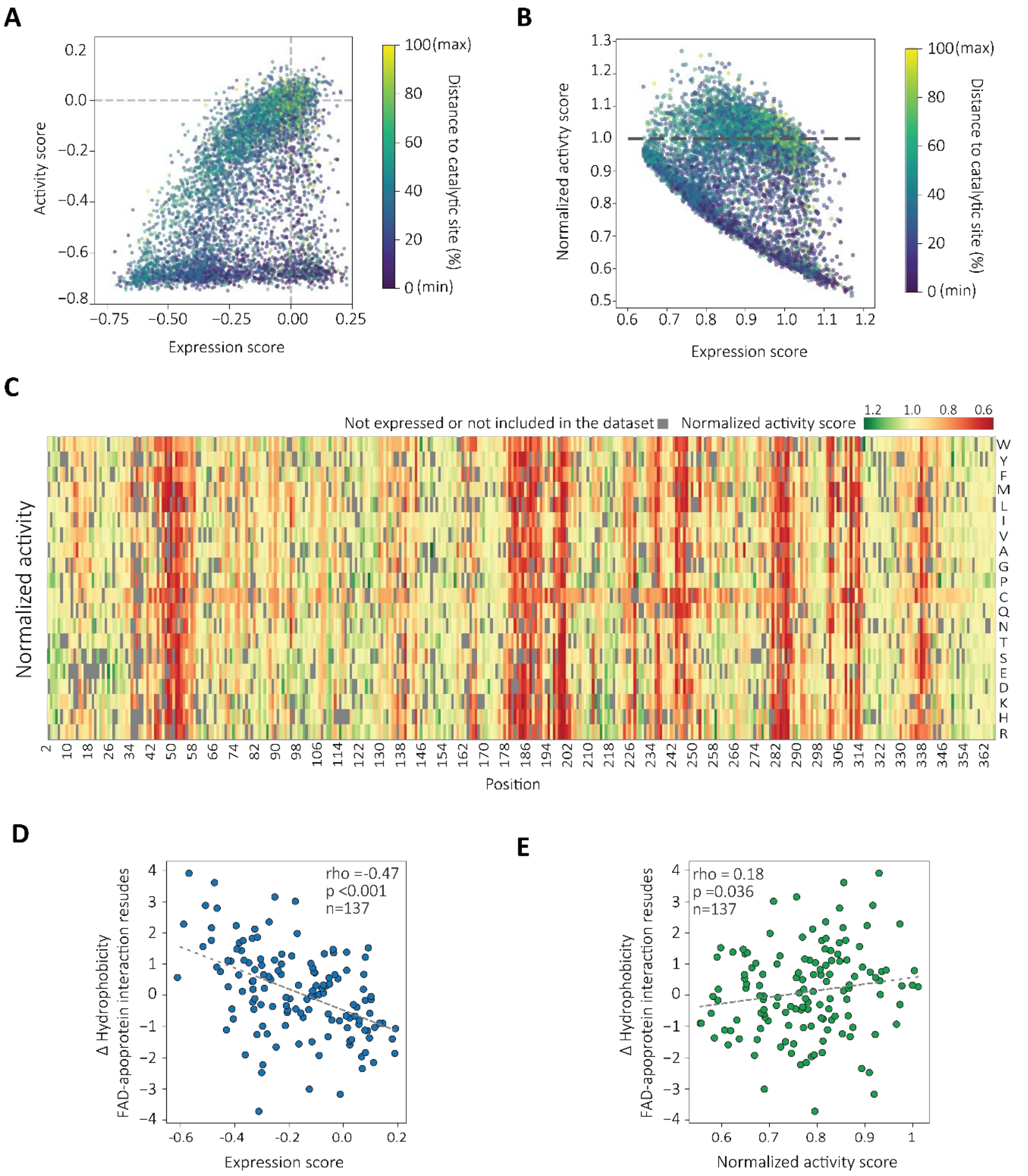

#### Figure S8. Deconvoluting activity from expression level

A) Scatter plot displaying the relationship between expression and activity fitness scores for 6,325 single mutant variants. B) Activity fitness scores were normalized over expression fitness scores (normalized activity) and visualized in relation to the expression score. Variants with normalized activity values greater than 1 (dashed line) represent single mutant variants where activity was improved independently of the expression level. C) Fitness heatmap of normalized activity scores along the sequence of DAOx. Gray boxes indicate mutant variants excluded from normalization due to their low expression scores (expression score <-0.65) and missing variants in the initial library. Interactive and color-blind accessible version of the heatmap can be found here:https://nash-lab.github.io/DAOx-DMS/heatmaps/hm_normalized_activity.html. D) Effect of changes in hydrophobicity at the core of DAOx on expression fitness. E) Effect of changes in hydrophobicity at the core of DAOx on normalized activity. Delta hydrophobicity values were calculated by subtracting the hydrophobicity score of the mutant residue from that of the wild-type residue for all variants with mutations at positions interacting with the FAD cofactor[34]. Conversely, positive delta hydrophobicity indicates a decrease in hydrophobicity whereas negative delta hydrophobicity indicates an increase in hydrophobicity.

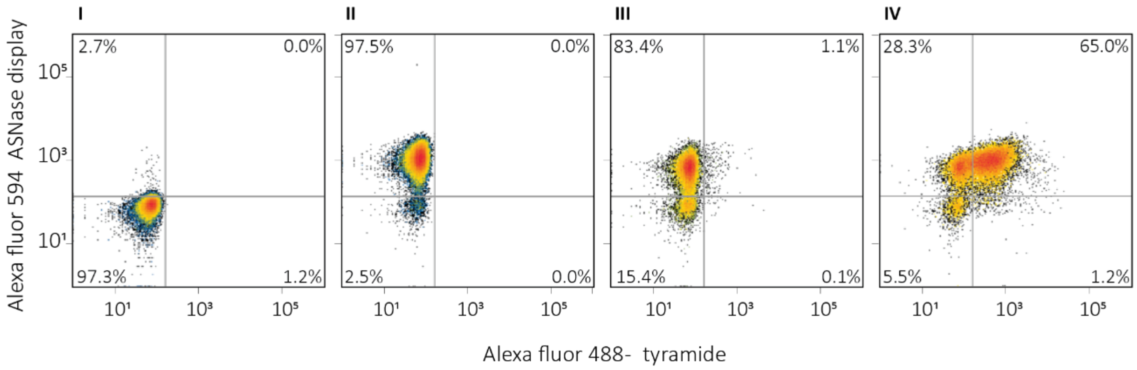

#### Figure S9. E.coli surface display and tyramide assay of Asparaginase 2 from *Erwinia chrysanthemi*

(I) *E.coli* (BL21 DE3) cells induced for the surface expression of *Erwinia chrysanthemi* Asparaginase 2 (ASNase) analyzed through flow cytometry. (II) *E.coli* surface display of ASNase detected through antibody staining and measured through flow cytometry. (III) Single cell tyramide activity assay negative control. A population of cells positive for the display of the ASNase was incubated with tyramide assay reaction mixture (1mM Asparagine, 10 µM *E. coli* Aspartate oxidase, 1/200 dilution of Alexa fluor™ 488 Tyramide, 50 µM FAD, 1.5% w/v alginate) lacking HRP for 45 minutes at room temperature. Cells were then analyzed through flow cytometry. (IV) Single cell tyramide activity assay. A population of cells displaying ASNase enzyme was assayed for the activity of the enzyme through a single cell tyramide assay. Cells were mixed with 1mM Asparagine, 10 µM *E. coli* Aspartate oxidase, 1 µM HRP, 1/200 dilution of of Alexa fluor™ 488 Tyramide, 50 µM FAD, 1.5% w/v alginate and incubated at room temperature for 45 minutes before being analyzed through flow cytometry.

#### Supplementary Notes

##### Note 1 - Cloning and yeast surface display of D-amino acid oxidase (DAOx).

To develop our workflow on DAOx, we codon optimized the cDNA sequence for *S. cerevisiae,* cloned it downstream of the Aga2 display anchor in a pYD1 derived plasmid, and transformed into *S. cerevisiae* EBY100. The fusion construct further contained a 6x-Histidine tag at its C-terminal end for antibody labeling. Yeast cells were cultured in galactose rich buffered media at 20°C for 48h. We found pH 7 to be optimal for Aga2-DAOx surface display, with more than 60% of the yeast population expressing WT DAOx under these conditions (Fig. S1*A*). To confirm correct folding and assembly of dimeric DAOx on the yeast surface, we tested the activity of displayed WT DAOx on D-alanine using a coupled HRP/Amplex Red assay. We detected DAOx activity for 500,000 cells displaying WT DAOx, with no fluorescence detected for yeasts lacking the displayed enzyme (Fig. S1*B*). To saturate DAOx and detect the maximum reaction rate, we used D-alanine at 35 mM, which was >40-fold higher than the reported $K_m$ of 0.8 mM[65], and ~5-fold higher than the $K_m$ we measured both for yeast displayed (7.316 ± 0.2816 mM) and soluble WT DAOx (6.965 ± 0.4003 mM). We varied HRP concentration in the assay and verified that, at an HRP concentration of 5.6 µM, the oxidation of D-alanine by DAOx was significantly rate limiting (Fig. S1*B*).
Next, we developed the single cell tyramide/peroxidase proximity labeling assay to measure DAOX activity in a format that would be compatible with pooled screening and DMS[26,66,67]. We mixed yeast cells displaying DAOx with 130 mM D-alanine, a 1/200 dilution of fluorescent tyramide-488, and 56.5 µM HRP. DAOx activity on D-alanine generated one equivalent of $H_2O_2$, which served as a downstream substrate for HRP-mediated proximity labeling of the yeast surface by tyramide-488. We observed a strong increase in green fluorescent signal for DAOx-expressing cells, while non-expressing cells showed no such increase (Fig. S1*C*). The specificity of the reaction was confirmed by incubating WT DAOx yeast in a reaction mixture lacking HRP, and detecting a more than 200-fold lower median fluorescence (Fig. S1*D*). We found that tyramide fluorescence of cell population was correlated with red fluorescence (expression stain) in the same population, demonstrating a quantitative relationship between DAOx display levels and tyramide labeling intensity (Fig. S1*D* inset).

#### Note 2 - Library construction and barcoding.

To construct a library for DMS analysis, we used one-pot site saturation nicking mutagenesis over the entire coding region of DAOx (codons 2 to 365)[5860]. To overcome the read length limitation of Illumina sequencing, we barcoded the variant library with 15 nucleotide unique molecular identifiers (UMIs) such that each variant was linked to one or more UMIs. The link between UMIs and variants was established through long read Pacbio sequencing (Fig. S2*A*). We estimated the total size of our library to be approximately 200,000 variants and stored the information for each UMI and the corresponding DAOx variant in a look-up table. The final UMI barcoded mutant library included nearly all possible DAOx single missense mutations (6,530 in total, 94.4% of theoretical) (Fig. S2*B*). Each mutant was linked to at least one UMI, with an average of 9 unique UMIs linked to each individual amino acid missense variant (Fig. S2*C*). In addition, our look-up table contained over 50,000 unique mutants carrying between 2 and 5 nucleotide mutations (Table S1). In the remainder of this work, we excluded all higher order mutants from the analysis and report on analysis only of single mutants. We tested the surface display of the variant library (Fig. S3), and based on the best expression conditions for both the library and WT DAOx, we selected 48h, 20°C and pH 7 as the final expression protocol for the screening stage.

#### Note 3 - Impact of mutant residue identity on the DAOx expression and enzymatic activity

Using data from the 6,399 single missense variants of DAOx for which both expression and activity scores were available, we grouped variants based on identity of the mutant residue and calculated their average effect on expression and activity fitness (Fig. 3A, B, Table S4). We plotted the percent fitness change relative to the average of all missense variants (Avg. Exp=-0.180, Avg. Act=-0.336). This analysis showed that proline (P) insertion had the most deleterious effect on both expression and activity, with average expression and activity scores ~57% (Exp=-0.281, n=328; Act=-0.527, n=328) lower than the average fitness of all variants in the respective screen. Cysteine (C) had a much less deleterious effect (~ +19% of average) on both properties (Exp=-0.148, Act= -0.263, n=331). We further found that neutral polar residues had a less deleterious effect on expression and activity fitness than the average mutation (Exp=-0.154 ,~ +14% Exp; Act=-0.299, ~ +10%; n=1287). Among non-polar residues, glycine (G) was the only substitution with lower than average fitness in both screens (Exp=-0.190, -6%; Act= -0.345, -3%; n=321). All other non-polar residues (leucine(L), valine (V), methionine (M), alanine (A) and isoleucine (I)) had an overall positive influence on expression and activity, with an average expression score ~17% and activity score of ~18% higher than the average of all missense mutations (Exp=-0.149; Act=-0.275). Insertion of charged amino acids less than histidine (H) had negative effects on catalytic activity with a fitness score ~12% lower than the average (Act=-0.379). Among charged residues glutamic acid (E) substitutions resulted in higher than average expression of +11% (Exp=-0.159; n=308) despite the general trend of charged amino acids having a negative impact on expression (Exp=-0.1900, -7%, n=1590). Among hydrophobic residues, bulky tryptophan (W) impaired both expression and activity fitness (Exp=-0.232, -29%; Act=-0.393, -17%; 332 variants) (Fig. 3*A* , Table S4).

#### Note 4 - Impact of the 3D structure on the expression and activity of DAOx

We next investigated whether the negative effects of proline insertion on DAOx expression and activity might be related to its ability to disrupt α helix and β sheet secondary structures due to the missing amide hydrogen necessary for hydrogen bonding. We found that inserting proline into α helix and β sheet regions of the DAOx significantly impaired both the expression and activity fitness compared to the effect of any of the other 19 amino acids when inserted in the same regions (Expression:α helix, Mann–Whitney U=85824.0, $p<0.001$, β sheet, Mann–Whitney U=44911.0, $p<0.001$; Activity: α helix, Mann–Whitney U=26334.0, $p<0.001$, β sheet, Mann–Whitney U=13744.0, p=1.24e-39, Fig. S5 *A*,*B* ). We expected regions of the protein possessing different secondary structures to have different tolerance levels for mutation that would be reflected in the expression and activity scores. We evaluated the average expression and activity fitness as a function of secondary structure of the mutated position and observed that linker regions had greater tolerance to mutations in both expression (avg. Exp=-0.129) and activity (avg. Act=-0.286) screens compared to structured regions (α helix, avg. Exp=-0.208, avg. Act=-0.348; β sheet, avg. Exp=-0.210, avg. Act=-0.396). β sheet regions had consistently the lowest tolerance to mutations (Fig. S5*C*,*D*).

#### Note 5 - Impact of wild type amino acid substitution on the DAOx expression and enzymatic activity

We analyzed expression and activity fitness as a function of the substituted wild type amino acids (Fig. S5*E,F*) and found that mutating low abundance hydrophobic core aromatic residues such as tryptophan (W) and tyrosine (Y) together with valine (V), one of the most abundant residues in DAOx (n=27), had the highest impact on the expression of the enzyme (avg. Exp V,Y,W=-0.276; avg. Exp all=-0.182) (Fig. S5*E*). Catalytic activity was strongly negatively affected by substitution of tryptophan (W) (avg. Act W=-0.593; avg. Act all= -0.342) (Fig. S5*F*). Mutation of polar or charged amino acids was comparatively less deleterious and was associated with on average higher scores (avg. Exp=-0.136, +25%; avg. Act,-0.260, +23%) than the other classes of amino acids (avg. Exp= -0.220; avg. Act=-0.410, Fig. S5*E,F*).
